# *De novo* determination of mosquitocidal Cry11Aa and Cry11Ba structures from naturally-occurring nanocrystals

**DOI:** 10.1101/2021.12.15.472578

**Authors:** Guillaume Tetreau, Michael R. Sawaya, Elke De Zitter, Elena A. Andreeva, Anne-Sophie Banneville, Natalie Schibrowsky, Nicolas Coquelle, Aaron S. Brewster, Marie Luise Grünbein, Gabriela Nass Kovacs, Mark S. Hunter, Marco Kloos, Raymond G. Sierra, Giorgio Schiro, Pei Qiao, Myriam Stricker, Dennis Bideshi, Iris D. Young, Ninon Zala, Sylvain Engilberge, Alexander Gorel, Luca Signor, Jean-Marie Teulon, Mario Hilpert, Lutz Foucar, Johan Bielecki, Richard Bean, Raphael de Wijn, Tokushi Sato, Henry Kirkwood, Romain Letrun, Alexander Batyuk, Irina Snigireva, Daphna Fenel, Robin Schubert, Ethan J. Canfield, Mario M. Alba, Frédéric Laporte, Laurence Després, Maria Bacia, Amandine Roux, Christian Chapelle, François Riobé, Olivier Maury, Wai Li Ling, Sébastien Boutet, Adrian Mancuso, Irina Gutsche, Eric Girard, Thomas R. M. Barends, Jean-Luc Pellequer, Hyun-Woo Park, Arthur D. Laganowsky, Jose Rodriguez, Manfred Burghammer, Robert L. Shoeman, R. Bruce Doak, Martin Weik, Nicholas K. Sauter, Brian Federici, Duilio Cascio, Ilme Schlichting, Jacques-Philippe Colletier

## Abstract

Cry11Aa and Cry11Ba are the two most potent toxins produced by mosquitocidal *Bacillus thuringiensis* subsp. *israelensis* and *jegathesan*, respectively. The toxins naturally crystallize within the host; however, the crystals are too small for structure determination at synchrotron sources. Therefore, we applied serial femtosecond crystallography at X-ray free electron lasers to *in vivo*-grown nanocrystals of these toxins. The structure of Cry11Aa was determined *de novo* using the single-wavelength anomalous dispersion method, which in turn enabled the determination of the Cry11Ba structure by molecular replacement. The two structures reveal a new pattern for *in vivo* crystallization of Cry toxins, whereby each of their three domains packs with a symmetrically identical domain, and a cleavable crystal packing motif is located within the protoxin rather than at the termini. The diversity of *in vivo* crystallization patterns suggests explanations for their varied levels of toxicity and rational approaches to improve these toxins for mosquito control.

## Introduction

The most commonly used biological insecticide for controlling mosquito vector populations is produced by the bacterium *Bacillus thuringiensis* subsp. *israelensis* (*Bti*) ^1^. Its highly potent mosquitocidal activity is due to three nanocrystalline forms of four protoxins, viz. Cyt1Aa, Cry11Aa, and co-crystallized Cry4Aa and Cry4Ba. These are produced during sporulation and are remarkably stable in a variety of conditions, but dissolve after ingestion under the high alkaline pH levels characteristic of the larval mosquito midgut ^2^. Solubilized protoxins are activated by insect gut proteases enabling binding to gut cell membranes, subsequent oligomerization, and ultimately gut cell lysis leading to larval death ^2^. *Bti* toxins are environmentally safe because they are much more specific for target mosquitoes than broad-spectrum chemical larvicides.

The most potent of the four *Bti* toxins is Cry11Aa, but it is poorly understood, in large part because unlike Cry4Aa, Cry4Ba, and Cyt1Aa, its structure is unknown. A related toxin produced by *Bt* subsp. *jegathesan* (*Btj*) is Cry11Ba, which is seven to thirty-seven times more toxic than Cry11Aa against major mosquito vector species belonging to the genera *Aedes, Anopheles*, and *Culex* ^3^, and in some bacterial hosts appears to form slightly larger crystals. Cry11Ba’s structure is also unknown, although it has been used in recombinant strains of *Bti* to improve mosquitocidal activity significantly ^3,4^. Thus, our goal was to determine the structures of Cry11Aa and Cry11Ba protoxins to help understand their mechanisms of crystallization that result in environmental stability and could possibly yield structural insights for increasing the efficacy of these proteins for mosquito control. Structure determination of Cry11Aa and Cry11Ba protoxins from natural nanocrystals requires cutting-edge technology. Conventional crystallography is limited to projects in which crystals are sufficiently large to mount and oscillate individually in a synchrotron X-ray beam. In the past, crystals of activated Cry4Aa ^5^, Cry4Ba ^6^ and Cyt1Aa ^7^ attained sufficient size by growing these *in vitro* from toxins dissolved from natural nanocrystals and activating the toxins enzymatically. However, Cry11Aa and Cry11Ba do not recrystallize *in vitro* from dissolved nanocrystals ^8^. Moreover, enzymatic activation is unwanted since our goal is to understand the pH-controlled mechanism of natural crystal dissolution. To observe the protoxin state in natural nanocrystals produced in bacterial cells, we applied serial femtosecond crystallography (SFX) at X-ray free electron lasers (XFEL) ^9–11^. In the SFX experiment, high brilliance XFEL beam pulses, each lasting only ~10-50 fs, intercept a series of nanocrystals, one pulse-per-crystal, eliciting the strongest possible diffraction signal from each tiny crystal before it vaporizes, and producing a series of diffraction snapshots, later assembled into a full data set. Feasibility of this strategy had been demonstrated by the recent elucidation of the full bioactivation cascade of Cyt1Aa ^12^.

Our success in determining the structures of Cry11Aa and Cry11Ba protoxins highlights the capability of XFEL sources to overcome limits of small crystal size. We relied on *de novo* phasing of the native SFX data because all attempts at molecular replacement (MR) failed despite detectable sequence similarity with ten structurally-determined members of the three-domain Cry δ-endotoxin family ^13–15^. We opted to derivatize our Cry11Aa nanocrystals with a recently-introduced phasing-agent, a caged-terbium compound, Tb-Xo4 ^16,17^. The phases obtained from single wavelength anomalous dispersion (SAD) were sufficient to reveal the Cry11Aa protoxin structure at 2.6 Å resolution and subsequently enable phasing of the Cry11Ba protoxin structure at 2.3 Å resolution by molecular replacement. In hindsight, we attribute the failure of early MR attempts to three extra β-strands in domain II which alter the relative orientation of the three domains in Cry11 toxins.

Our studies of Cry11Aa and Cry11Ba crystals reveal a new paradigm of molecular packing among Cry δ-endotoxins reported thus far. In particular, the cleavable peptides that constitute important crystal contacts are located near the middle of the toxin sequence, rather than at the termini. Molecules pack in tetramer units, exhibiting D2 symmetry; these tetramers in turn pack in a body centered pattern (like a 3-dimensional brick-wall in which successive rows are offset by half a brick). To achieve this pattern, each of the three domains in a Cry11 molecule packs with an identical domain from a symmetry related molecule: domain I packs with domain I, II with II, and III with III. Thus, each Cry11 domain fulfills two biological roles: a dimer interface manifested in the crystalline state, and a functional role manifested in the soluble state: target recognition (domain II), oligomerization (domain III) and pore formation (domain I) ^18^. Differences in the size and composition of the three packing interfaces explains shape and size differences between Cry11Aa and Cry11Ba nanocrystals. Structure-guided site-directed mutagenesis verifies which residues affect crystal size, pH sensitivity of the crystal, and toxin folding. Our results elucidate the Cry11Aa and Cry11Ba bioactivation cascade and enable development of new, rational strategies for improved mosquito control.

## Results

### De novo phasing of Cry11Aa and Cry11Ba structures by SFX

*In vivo*-grown crystals of Cry11Aa and Cry11Ba protoxins exhibit distinct morphologies, which initially concealed a surprising conservation of their crystal packing patterns. Cry11Aa crystallizes as hexagonal plates and Cry11Ba crystallizes as larger bipyramidal crystals (Fig. 1 a,b) as reported earlier ^4^. These morphological distinctions cannot be attributed to differences in crystallization mechanisms in their parent organisms, *Bti* and *Btj*, since both protoxins were recombinantly produced in the same host organism, an acrystalliferous strain of *Bti* (4Q7). Cry11Aa and Cry11Ba protoxins are expected to share structural resemblance to each other since the two sequences share 54% identity; however, 46% non-identity at the molecular level could easily produce large differences at the macroscopic level of crystal morphology. Moreover, the sequence of Cry11Ba is extended by 77-residues at its C-terminus, potentially also affecting differences in crystal packing (Supplementary Fig. 1). Interestingly, this extension has been identified as a low complexity region (LCR) by both CAST ^19^ and SEG ^20^ computational methods, which implicates the extension in the mechanism of crystal nucleation. At this point in our studies, the balance of evidence suggested that sequence divergence was likely to have erased the crystal packing pattern that early ancestors of today’s Cry11Aa and Cry11Ba presumably once shared.

**Fig. 1.**
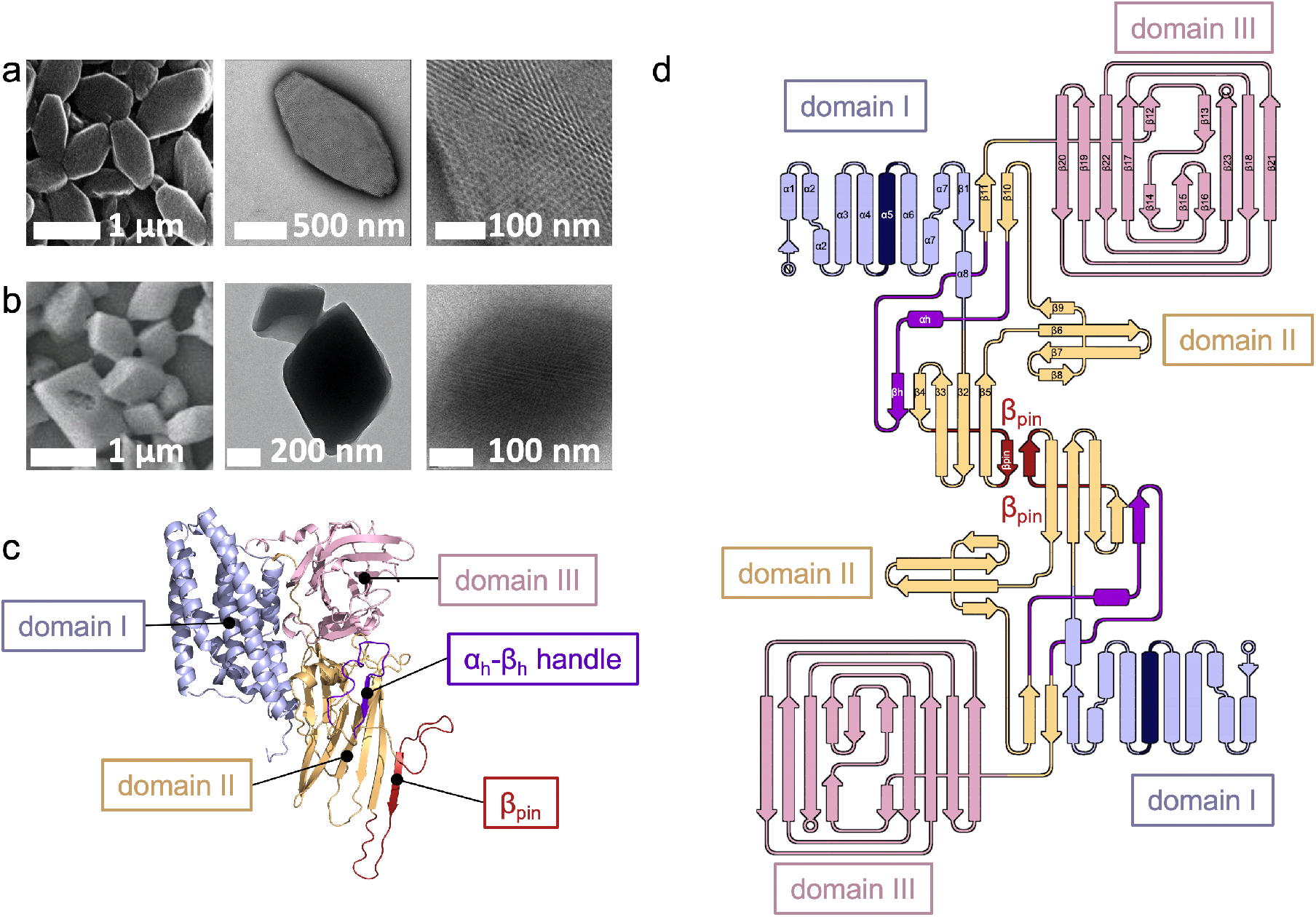
Crystals and overall fold of Cry11 toxins. **a-b,** scanning (left; SEM) and transmission (middle, right; TEM) electron micrographs of gold plated and negatively-stained Cry11Aa (**a**) and Cry11Ba (**b**) crystals, respectively. The right panels show a close-up view of the crystal surface. **c,** Cry11Aa crystal structure, depicted as a cartoon. Domain I is shown in blue; domain II is shown in orange except for the α_h_β_h_-handle and β_pin_ which are shown in purple and red, respectively; domain III is shown in pink. **d**, Topology diagram of a Cry11Aa dimer. Domain I is shown in green, except for central helix α5, which is shown in blue: domain II is shown in magenta, except for the α_h_β_h_-handle, which is shown in purple; and domain III is shown in cyan, respectively. The two monomers in a dimer assemble via the β_pin_, resulting in the formation of a large β-sheet.

Our diffraction experiments yielded the first hint that Cry11Aa and Cry11Ba shared a conserved crystal packing pattern. We collected diffraction data from Cry11Aa and Cry11Ba nanocrystals injected in the vacuum chamber of the CXI-SC3 micro-focused beamline at the Stanford Linear Accelerator Center (SLAC) Linac Coherent Light Source (LCLS) ^21^ using a microfluidic electrokinetic sample holder (MESH) ^22^ (Cry11Ba crystals) or a gas-dynamic virtual nozzle (GDVN) ^23^ (Cry11Aa crystals). The underlying similarity in the packing of Cry11Aa and Cry11 Ba became evident when their diffraction patterns were collected and indexed, revealing similarly sized unit cells (a~58; b~155; c~171 Å; α=β=ɣ=90°), albeit belonging to two different space groups: *I*222 and *P*2_1_2_1_2, respectively (Table 1). Conservation of unit cell parameters hinted that this crystal packing pattern is special, evolved to perform a function more intricate than just storing protein.

**Table 1.** Data collection and refinement statistics Cry11Aa and Cry11Ba.

|  | Cry11Aa<br>pH 7 | Cry11Aa-TBXO4<br>pH7 | Cry11Ba<br>pH 6.5 | Cry11Ba<br>pH 10.4 |
| --- | --- | --- | --- | --- |
| PDB ID | 7QX4 |  | 7QYD | 7R1E |
| **Data collection** |  |  |  |  |
| Space group | *I* 2 2 2 | *I* 2 2 2 | *P* 2<sub>1</sub> 2<sub>1</sub> 2 | *P* 2<sub>1</sub> 2<sub>1</sub> 2 |
| Cell dimensions (Å) | 57.64 ± 0.19 | 57.64 ± 0.15 | 168.18 ± 0.19 | 167.50 ± 0.29 |
|  | 155.69 ± 0.80 | 156.29 ± 0.73 | 158.45 ± 0.26 | 157.99 ± 0.47 |
|  | 171.14 ± 0.54 | 170.75 ± 0.40 | 57.51 ± 0.08 | 57.43 ± 0.14 |
| Wavelength (Å) | 1.27 | 1.27 | 1.30 | 1.30 |
| X-ray beam focus (μm) | 5 | 5 | 1 | 1 |
| No. collected frames | 792623 | 558747 | 813133 | 990643 |
| No. indexed frames | 48652 | 77373 | 19708 | 15689 |
| No. merged crystals | 50613 | 88511 | 19708 | 15689 |
| Resolution range (Å) | 33.55 – 2.60 | 33.51 – 2.55 | 44.73 – 2.20 | 35.72 – 2.55 |
|  | (2.66 – 2.60) | (2.61 – 2.55) | (2.24 – 2.20) | (2.59 – 2.55) |
| No. observations | 8253629 (365007) | 14069217 (640046) | 3996555 (31710) | 3747530 (52399) |
| No. unique reflections | 24198 (1583) | 48634 (3297) | 78897 (3815) | 50646 (2488) |
| <I/σ (I)> | 9.50 (1.16) | 11.23 (1.62) | 3.87 (0.50) | 3.65 (0.84) |
| R<sub>split</sub> (%) | 10.73 (95.40) | 7.97 (70.58) | 18.0 (111.8) | 24.50 (97.10) |
| CC<sub>1/2</sub> | 1.00 (0.38) | 1.00 (0.68) | 0.99 (0.12) | 0.984 (0.082) |
| Completeness (%) | 99.9 (100.0) | 100.0 (100.0) | 97.9 (80.9) | 99.4 (100.0) |
| Multiplicity | 341.09 (230.58) | 289.29 (194.13) | 50.66 (8.3) | 71.34 (21.01) |
| **Anomalous data** |  |  |  |  |
| Completeness (%) |  | 100.0 (100.0) |  |  |
| CCano |  | 0.26 (0.00) |  |  |
| CRDano |  | 1.35 (1.01) |  |  |
| **Refinement** |  |  |  |  |
| Resolution range (Å) | 33.55 – 2.60 |  | 44.77 – 2.20 | 35.72 – 2.55 |
| No. reflections | 24196 |  | 70858 | 50657 |
| R<sub>work</sub>/R<sub>free</sub>* | 17.2 / 24.1 |  | 18.9 / 23.8 | 23.8 / 19.2 |
| No. atoms |  |  |  |  |
| Protein | 5080 |  | 10083 | 9961 |
| Water | 261 |  | 623 | 98 |
| B-factors (Å<sup>2</sup>) |  |  |  |  |
| Main chain | 50.47 |  | 41.82 / 41.06<sup>§</sup> | 41.02 / 42.58 |
| Side chain | 51.44 |  | 46.11 / 45.55 | 43.26 / 44.59 |
| Water | 46.17 |  | 41.36 | 40.04 |
| R.m.s.d. |  |  |  |  |
| Bonds lengths (Å) | 0.004 |  | 0.008 | 0.009 |
| Bonds angles (°) | 0.633 |  | 1.324 | 1.590 |

\* R<sub>free</sub> is calculated using 5 and 10% % of random reflections excluded from refinement.<sup>§</sup> Average B-factor for chain A / chain B

To gain further insight into Cry11Aa and Cry11Ba crystal packing, we depended on *de novo* methods to solve the crystallographic phase problem. Initial attempts to acquire phases from homologous structures by molecular replacement (MR) failed, suggesting Cry11Aa and Cry11Ba contained novel features, not present in the PDB. Our search models included structures of Cry δ-endotoxins homologs (exhibiting up to 26% sequence identity to our two targets) and homology models produced using Robetta ^24^ (http://robetta.bakerlab.org/) and SwissProt ^25^ (https://www.ebi.ac.uk/uniprot/). After MR failed, we turned to *de novo* phasing methods. We soaked Cry11 nanocrystals with conventional heavy atom derivatives including gadolinium, gold, platinum, and mercury salts, but they failed to produce interpretable isomorphous or anomalous difference Patterson peaks. Finally, a recently introduced caged-terbium compound ^16,17^, Tb-Xo4, produced a successful derivative of Cry11Aa (after a 30h soak at 10 mM concentration), and phases were determined by the single wavelength anomalous dispersion (SAD) method at 2.6 Å resolution (using anomalous signal up to 3.5 Å). Two Tb-Xo4 molecules were identified bound to the single Cry11Aa molecule in the asymmetric unit (isomorphous peaks at 23 and 9 σ, and anomalous peaks at 33 and 8.1 σ, respectively; Supplementary Fig. 2a). The success of Tb-Xo4 can be partly ascribed to the dramatically high anomalous dispersion signal (*i.e*. f’ and f”) of terbium, but likely also stems from stronger binding of TbX04 to the protein owing to presence of an organic cage; indeed, f’ and f” of Gd and Tb are similar at the X-ray energy used for data collection (9 keV). Regardless, phases were of sufficient quality to reveal all Cry11Aa residues from N13 to the C-terminal K643.

The Cry11Ba structure was thereafter phased successfully by MR using the Cry11Aa structure as a search model. *A posteriori*, we discovered that two of the heavy atom compounds that we used for soaking actually did bind Cry11Ba (Supplementary Fig. 2b-c). Difference Fourier maps revealed 7-8 σ peaks indicating Pt bound near Met 19 and 200, and Gd bound near Asp83 and Asp427 (Supplementary Fig. 2b). Surprisingly, however, there were no peaks in the anomalous difference Fourier maps. We speculate that if we had achieved higher heavy-atom occupancy and/or higher multiplicity in our measurements, the anomalous signal would have been strong enough to detect and perhaps used for phasing. Our MR-phase 2.3 Å resolution map reveals two Cry11Ba molecules in the asymmetric unit. All residues are visible except for the N-terminus (residues M1-N16), two loops (residues G330-E340, and D352-I358) and the C-terminal extension (residues T654-K724). The lack of order in this extension is not surprising given the low complexity of its sequence.

### *Cry11 domain organization is similar to* δ-endotoxins, but exhibits some *non-canonical features*

Cry11Aa and Cry11Ba structures maintain the three-domain organization characteristic of Cry δ-endotoxins ^13,26^ (Fig. 1c, Supplementary Fig. 3). Domain I is implicated to form a pore in the target membrane. Like other Cry δ-endotoxins it forms a seven-α-helix bundle; at the center of the bundle is α5 (residues 146-170), surrounded by the remaining six helices. Domain II is implicated to recognize mosquito-specific receptors. It forms a β-prism composed of three-β-sheets, wherein the first two β-sheets (β4-β3-β2-β5 and β8-β7-β6-β9) each adopts a Greek-key topology while the third β-sheet is three-stranded (β1-β10-β11). Domain III is implicated to oligomerize. It forms a β-sandwich of two antiparallel five-stranded β-sheets (viz. β20 – β19 – β22 – β17 – β4 – ^β12^/_β14_ and β15 – ^β13^/_β16_ – β23 – β18 – β21) forming a jelly-roll topology, whereby ^β12^/_β14_ and ^β13^/_β16_ are interrupted β-strands contributed by two non-consecutive shorts β-strands, which appose and intercalate one after the other onto β4 and between β15 and β23, respectively (Figure 1d).

The closest homolog of known structure to Cry11 toxins is *Bt kurstaki* (*Btk*) Cry2Aa (PDBid: 1i5p), with a sequence identity of 26.6 and 23.6 % and main-chain rmsd of 3.7 and 4.0 Å, with respect to Cry11Aa and Cry11Ba, respectively. As with Cry2Aa, the Cry11Aa toxins feature a long insert (27 residues in Cry2Aa; 21 residues in the Cry11 toxins) between strands β10 and β11, which together with domain-I β1, form the third β-sheet of the domain-II β-prism. This insert, which features a short α-helix (α_h_) and a β-strand (β_h_), folds like a handle, and is therefore referred to as the α_h_β_h_-handle, throughout the manuscript (Fig. 1c, Supplementary Fig. 3). The α_h_β_h_-handle fastens domain II onto domain III through direct (e.g. in Cry11Aa, D443(OD2)-R502(NH2); D443(O)-R502(NH1); L447(N)-S503(O)) and water mediated H-bonds (T446(OG1)/T448(O)-Wat72(O)-R502(N); T448(OG1)/V499(O)-Wat308(O)-D501(OD1); T448(N)/L447(N)-Wat65(O)-S503(OG)/(O)), and enables the burying of domain-II α8 at an interface formed by α_h_β_h_, α6-α7 (domain I), β10-β11 (domain II), β15 and the β13-β14 and β15-β16 loops (domain III), and the α9 helix connecting domain II and domain III (D469-K478 in Cry11Aa) (Supplementary Fig. 4). The firm hold of α8 enables the three domains to be more tightly packed in Cry2Aa and Cry11 toxins than in other Cry toxins (e.g. *Btt* Cry3Aa or *Btk* Cry1Ac). Additionally, strand β_h_ lays aside strand β4 thereby expanding – and consequently, stabilizing – the first β-sheet of domain II (β_h_-β4-β3-β2-β5). Also, alike Cry2Aa, the Cry11 toxins feature a smaller β-prism due to deletions in the second constitutive β-sheet, namely between β7 and β8 (6 and 10 residues missing in Cry2Aa and Cry11 toxins, respectively), and between β9 and β10 (14 and 15 residues missing in Cry2Aa and Cry11 toxins, respectively; Supplementary Fig. 3). The Cry11 toxin structures are, however, specific in that a 36 to 38 residue insertion is observed between strands β4 and β5, contributing an additional β-strand to the first β-sheet of domain-II – thereafter referred to as the β_pin_ (Fig. 1c). As the β_pin_ lays along a two-fold axis, two large β_h_-β4-β3-β2-β_pin_ – β_pin_-β2-β3-β4-β_h_ sheet are formed between symmetry related dimers (AC or BD, interface #3), yielding the crystallizing tetramer (Fig. 2b, e). We noted earlier that the BSA at the tetramerization interface is 33% lower in Cry11Ba, pointing to higher flexibility; this hypothesis is supported by the absence of interpretable electron density for residues at the N-terminus (330-340) and C-terminus (352-360) of the β_pin_ in the Cry11Ba structure. Also noteworthy is that Cry11 toxins feature a conserved N/D-DDLGITT insertion between β21 and β22, and deletions (>3 residues) between α3 and α3 (−5 and −8 residues with respect to *Btk* Cry2Aa and *Bt tenebrionis* (*Btt*) Cry3Aa), and β20 and β21 (−10 and −9 residues with respect to *Btk* Cry2Aa and *Btt* Cry3Aa). Altogether, these changes render Cry11 toxins uniquely large from the structural standpoint, with predicted radii of gyration of 27.5 and 26.7 Å for Cry11Aa and Cry11Ba, compared to 25.0 and 25.6 Å for *Btk* Cry2Aaa and *Btt* Cry3aa, respectively.

**Fig. 2.**
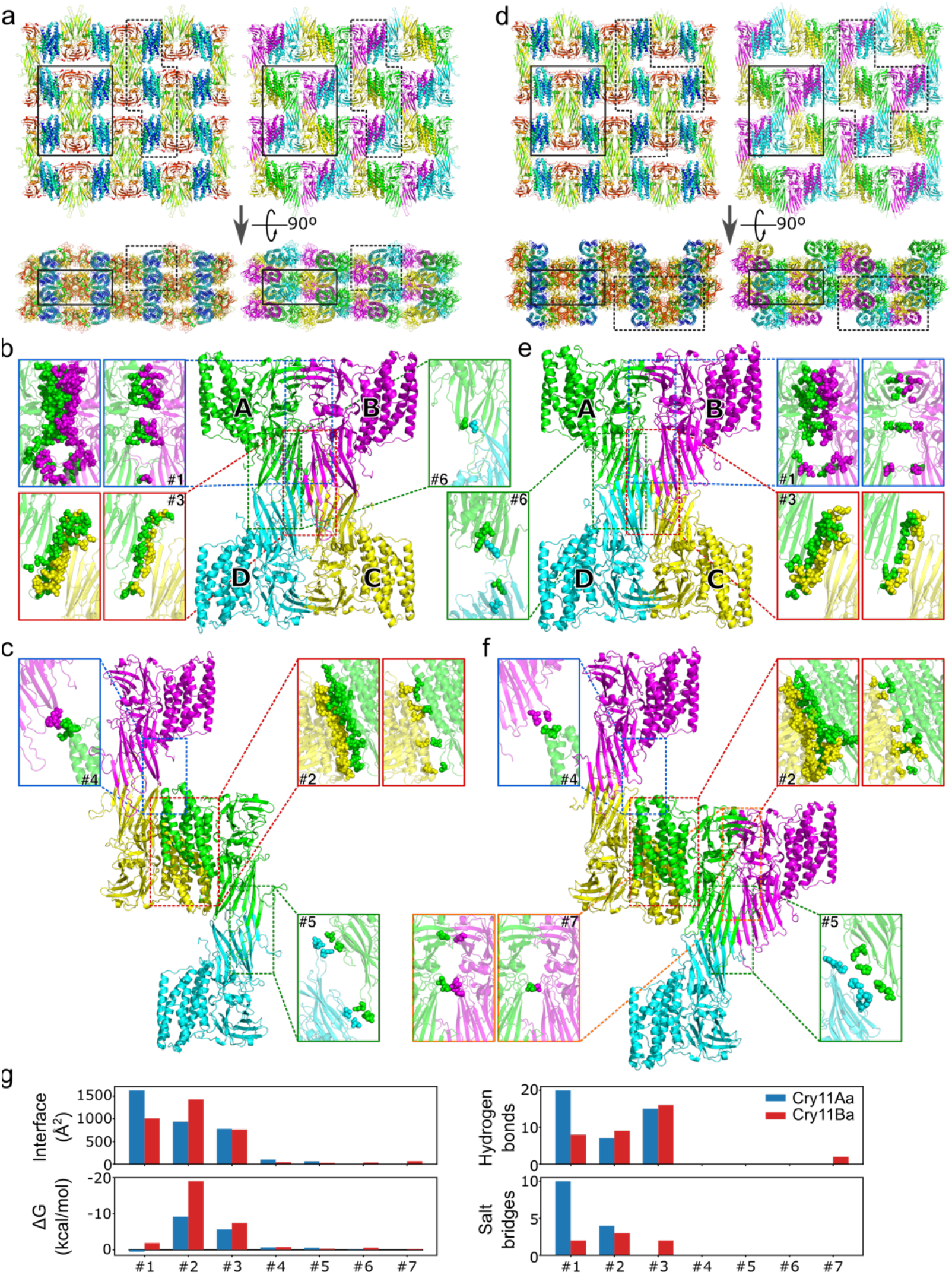
Monomer interactions in Cry11Aa and Cry11Ba. **a,** Cry11Aa crystal packing, coloured according to sequence (from blue to red) indicating the domain-based assembly; and coloured according to tetramer assembly (see panel (**b**)). The highlighted areas indicate the regions shown in (**b**) (full line) and (**c**) (dashed line). **b,** Cry11Aa tetramer with zoom on each of the three interfaces identified by PISA (interface #1, #3 and #6), with the involved residues depicted as spheres. For interfaces with hydrogen and/or salt bridges (see g), an additional (right) image shows only those residues that make up these interactions. **c,** Cry11Aa crystal assembly by interactions between neighbouring tetramers, formed by interface #2, #4 and #5, visualized as in b. **d,** Cry11Ba crystal packing, coloured as in (**a**). **e,** Cry11Ba tetramer with zoom on the interfaces as in (**b**). **f,** Cry11Ba crystal assembly, visualized as in (**c**). As compared to Cry11Aa, Cry11Ba crystals contain an additional interface #7 between an A-B pair from two neighbouring tetramers. **g,** interface statistics as identified by PISA for Cry11Aa (blue) and Cry11Ba (red).

### All domains engage in producing the in vivo crystal lattice

Examination of packing interfaces reveal that all three domains are involved in the formation and stabilization of Cry11Aa and Cry11Ba nanocrystals. The *in vivo* crystallization pathway can be best trailed from Cry11Aa crystals, which feature a single monomer per asymmetric unit and build on six packing interfaces burying a cumulated surface area (BSA) of 3514.5 Å^2^, corresponding to 13.1 % of the total protein area. The main building block of Cry11Aa crystals consists of a tetramer with a total BSA of 9663.0 Å^2^ and a predicted binding energy of −12.5 kcal.mol^-1^ at pH 7 by PISA ^27^ (Fig. 2a,b). Supported by two of the six packing interfaces, the tetramer builds from the cross-association (AC or BD; interface #3) of two dimers (AB or CD) along the 2-fold axis contributed by domain II (Fig. 2b). Each dimer is itself composed of monomers associated along another 2-fold axis contributed by domain III and by strand β4 and the β10-β11 hairpin (P433-P457) in domain II. The tetramer is further stabilized by a minor interface between apices from domain II (interfaces AD or BC; interface #6). Crystals grow from the piling in a honey-comb brick-wall fashion of such tetramers, as a result of a face-to-back interaction between domains I from symmetry-related molecules (interface #2; Fig. 2c). Cry11Aa crystals are further cemented by two additional minor interfaces. The first involves the apex of the second β-sheet of domain II (interface #5) from facing monomers in each dimer (AD or BC) of the stable tetramer. The second occurs between the α3-α4 loop of domain I in one tetramer and the apex of the second β-sheet of domain II in another tetramer (interface #4).

The similarity between the packing of Cry11Aa and Cry11Ba crystals makes it reasonable to propose that the latter also forms from the assembly of tetramers (Fig. 2d, e, f), despite failure of PISA to identify the tetramer as a (meta-)stable building block for Cry11Ba crystals. In these, two molecules are found in the asymmetric unit, associated through the face-to-back interface between domains described above for Cry11Aa monomers (interface #2; Fig. 2f). The BSA at this interface is 1429.0 Å^2^, *i.e*. 53% higher than in Cry11Aa (Fig. 2g). However, BSAs at the interface formed by domains III and II, which associates monomers into a dimer is 38% lower than the homologous interfaces in Cry11Aa (Fig. 2e, g). Thus, the intermolecular contact between piled tetramers is larger in the Cry11Ba crystals, despite an overall looser packing of monomers in the crystals, with an average BSA at crystal contacts of 3385.1 Å^2^ per monomer, corresponding to 12.6 % of the total protein area. The increased BSA between tetramers (contributed by the large face-to-back interface between domains I), and the presumably higher flexibility in the Cry11Ba tetramers, could be at the origin of Cry11Ba packing into larger three-dimensional crystals. Regardless, our structures evidence that each domain exhibits a dual role in Cry11 toxins, namely the formation and stabilization of *in vivo*-grown nanocrystals, and execution of a domain specific function. The latter comprises pore formation (domain I), receptor-recognition and membrane-insertion (domain II), and oligomerization and stabilization of the toxic pore conformation (domain III) ^26^.

### Drastic conformational changes drive crystal dissolution

We sought to characterize the conformational changes that ensue pH elevation, preceding dissolution of the crystals in the mosquito larvae gut ^28^. As the crystals are naturally labile at pH 11, we aimed at collecting data from crystals soaked at a lower pH, hypothesizing that early conformational changes would show but the crystal packing still hold. In the case of Cry11Aa crystals, diffraction quality was decreased dramatically at pH values of 9.5 (CAPS buffer, glycerol 30%) and above, preventing collection of a sufficiently large number of diffraction patterns to produce a high-pH dataset. Hence, large conformational changes occur in Cry11Aa at pH as low as 9.5, opposing diffraction quality, despite crystals dissolving as of pH 11 only (Fig. 3a). In the case of Cry11Ba, ~3 Å diffraction was preserved up to pH 10.4 (Table 1). Comparison between the refined ‘pH10.4’ and ‘pH6.5’ structures points to large inter-domain rearrangements induced by pH increase. Detailed analysis of structural changes at the side chain level was yet prevented by the limited resolution of the ‘pH10.4’ dataset. A 1% unit cell contraction, and hence tighter crystal packing, was observed in the ‘pH10.4’ crystals in comparison to the pH 6.5 crystals. However, because a higher glycerol concentration was used for injection of Cry11Ba crystals at pH 10.4, we cannot exclude that unit cell contraction might be caused by crystal dehydration.

**Fig. 3.**
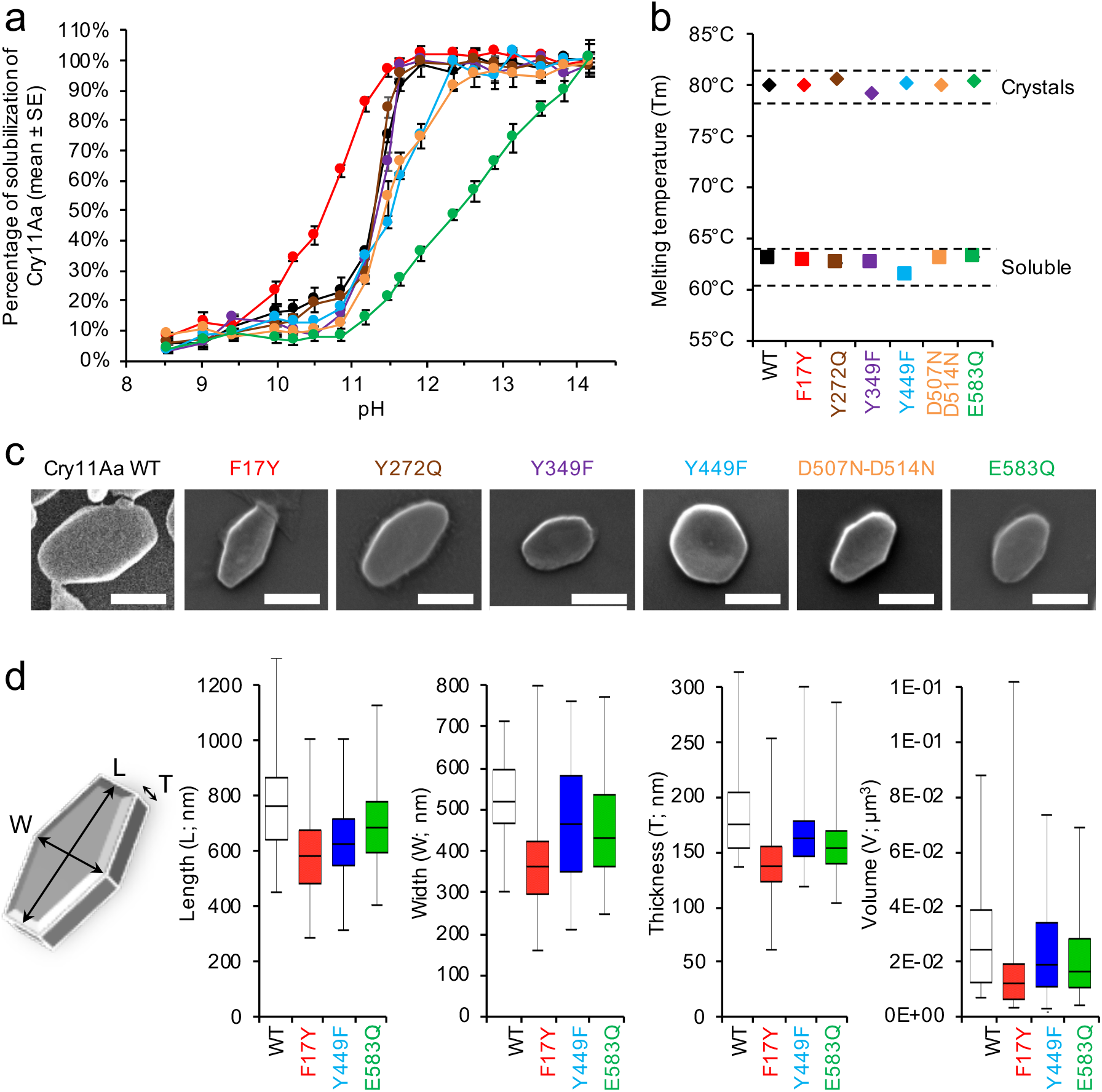
Point-mutations of Cry11Aa affect the shape, size and pH-sensitivity of *in vivo-* grown nanocrystals. **a, C**rystals from mutants exhibit similar sigmoidal patterns of crystal solubilization as a function of pH, except F17Y and E583Q that are more and less sensitive to pH, respectively. Error bars indicate the standard error of the measurements. **b,** Cry11Aa WT and mutants exhibit similar heat stability. As expected, toxins are more stable (+ 17.5 ± 0.3°C) in their crystalline than soluble form, irrespective of the mutation. **c,** Visualization of a representative crystal for Cry11Aa WT (black) and mutants F17Y (red), Y272Q (brown), Y349F (purple), Y449F (blue), D507N-D514N (orange) and E583Q (green) by SEM (scale bar = 500 nm). **d,** Crystals of Y449F, F17Y and E583Q imaged by AFM were all smaller in length (L), width (W), thickness (T) and volume than WT highlighting a perturbation of the intrinsic crystal organization induced by these mutations. In each graph, the boxes represent the lower and upper quartiles around the median. The whiskers indicate the minimum and maximum values.

### Crystals are made of full-sized monomers of Cry11 toxins

In both Cry11Aa and Cry11Ba toxins, the β_pin_ (residues E339-Q350 and I341-Y350, respectively) is a ~10-residue long β-strand that hydrogen-bonds with a two-fold related symmetry mate, contributing the interface that assembles dimers (AC and BD) into stable tetramers. This strand is bordered on each side by the only two loops that have disordered electron density in Cry11Ba (missing residues G330-E340 and D352-I358) and are comparatively difficult to interpret in Cry11Aa (F330-D334 and Q350-E355), respectively. As Cry11Aa N335-Y349 and Cry11Ba I341-N351 regions match the enzymatic cleavage site known to generate the two activated fragments of ~32 and ~36 kDa ^29,30^ upon proteolytic activation in the mosquito larvae gut, we asked whether disorder in the F330-D334 (G330-E340) and Q350-E355 (D352-I358) loops serves the purpose of enabling facilitated access of proteases to Cry11Aa (Cry11Ba) cleavage sites or if each monomer occurs in natural crystals as two polypeptide chains cleaved prior or during crystal formation. SDS PAGE analysis of Cry11Aa (12% gels, heating at 95°C for 5 min, presence of DTT and SDS; Supplementary Fig. 5) resulted in a major band ~70 kDa, in line with previous reports ^31–33^. As the denaturing treatment would have broken any disulfide-bridge or non-covalent interactions that could maintain cleaved fragments together, this result suggests that Cry11Aa occurs in crystals as a full monomer. We further verified this hypothesis by use of MALDI TOF mass spectrometry. In MALDI mass spectra collected after direct solubilization of the natural crystals in sinapinic acid matrix in presence or absence of DTT, we observed main peaks at *m/z* of 72246 and 72235 (mass error: ± 100 Da) and 36154 and 36129 Da, respectively, in agreement with expected molecular masses for singly- and doubly-charged ions of a full-size monomer (expected mass: 72.349 kDa) (Uniprot accession number: P21256; Supplementary Fig. 6). However, because proteolytic activation is as well expected to yield a 36 kDa fragment, in addition to a 32 kDa fragment for which a minor peak was present in the MALDI-TOF mass spectra, we resorted to native mass spectrometry to assert that the ~72.240 and ~36.140 kDa peaks originated from the same species – rather than being indicative of the crystallization of proteolytic products. With this approach, we could confirm that upon dissolution of Cry11Aa crystals, a 72.345 kDa fragment is released, corresponding to the full-size monomer (Supplementary Fig. 7a). Moreover, both incubation of solubilized toxin at room temperature (RT) for 2 h (Supplementary Fig. 7b) and use of increased collision energy (Supplementary Fig. 7c, d) failed at yielding a signature for the two polypeptides that would have been generated if cleavage at position 329 had occurred. We conclude that natural crystals of Cry11Aa, and possibly Cry11Ba, grow from the addition of full-size monomers, and that disorder in the F330-D334 (G330-E340) and Q350-E355 (D352-I358) loops could serve the purpose of enabling facilitated access of proteases to Cry11Aa (Cry11Ba) cleavage sites. Considering proteinase K as a surrogate analogue for mosquito larvae gut proteases ^34^, one would expect the β_pin_ to be released upon proteolytic activation, suggesting that the role of the latter is to promote *in vivo* crystallization. We note that other cleavage sites are predicted, which would release the first six residues and last two β-strands (β22-β23), as well as rescind the covalent association between domain I and domains II and III, thereby leaving non-covalent interactions surfaces as the sole glue between them.

### Mutagenesis to alter crystal formation and dissolution

We proposed earlier that the packing of Cry11Ba into slightly larger crystals than Cry11Aa could stem from differences in the extent and nature of the interfaces which support dimerization, tetramerization and piling of tetramers into crystals (Fig. 2). Considering recent evidence linking LCR regions with diverse functions including chaperoning ^35^ and reversible oligomerization, we further asked whether or not presence of the 77-residue LCR region at the C-terminus of Cry11Ba plays a complementary role in the promotion of crystal formation. A chimera was therefore designed, coined C11AB, wherein the LCR region of Cry11Ba was fused to the C-terminal end of Cry11Aa (Material and Methods; Supplementary Fig. 8a). C11AB was produced at the expected size but at a lower yield than Cry11Aa WT (Supplementary Fig. 8b). Atomic force micrographs (AFM) revealed the presence of multiple needle-like inclusions in the parasporal envelope encasing the crystals, suggesting that presence of Cry11Ba-LCR at the C-terminal end of Cry11Aa favors nucleation, but not crystal growth (Supplementary Fig. 8c).

Seven Cry11Aa mutants were additionally designed with the aim to probe the involvement of Cry11Aa intra- and inter-molecular interfaces in toxin stability, crystal formation and dissolution. Each mutant was designed to challenge a specific interface and served as a coarse proxy to evaluate its pH sensitivity and putative participation in the crystal dissolution mechanism. First, we asked whether the intra-chain stabilization of α8 at an interface contributed by the three domains (namely, α_h_β_h_, α6-α7, α9, β10, β11, β15 and the β13-β14 and β15-β16 loops) could play a role in crystal dissolution. Residues central to this interface are Y272, D514 and D507, which H-bond to one another and to Y203, R222, T249, S251 through direct and water-mediated interactions (W253 and W267), connecting the three domains (Supplementary Fig. 9a, Supplementary Table 1a). Upon pH elevation, Y272, D514 and D507 are all expected to be deprotonated, which should result in electrostatic repulsion and thence dissociation of the three domains. To test the hypothesis, we produced three Cry11Aa mutants intended to eliminate pH sensitivity of the above-described H-bonds. Neither did the Y272Q nor D507N-D514N mutations impact the overall stability of the toxin, in the soluble or crystalline form (Fig. 3b), but their combination in the triple Y272Q-D507N-D514N mutant resulted in an unexpected abolishment of the ability of Cry11Aa to form crystals *in vivo* – possibly due to improper folding (Supplementary Fig. 10). The Y272Q mutation had no effect on the pH sensitivity of Cry11Aa crystals, while only a minor effect was seen with the D507N+D514N mutant (Fig. 3a). Thus, pH-induced deprotonation of amino acids involved in the stabilization of α8 at the interface between the three domains does not play a role in the initial steps of crystal dissolution, possibly because of their deep burial at the interface. We note that the above-mentioned residues and their interactions are all strictly conserved in Cry11Ba (viz. Y273, D518, D511, Y203, R222, T249, S251, W253 and W268).

We then focused on Cry11Aa E583, a residue sitting at the intramolecular interface between domain I and domain III. This β21 residue, condemned to be anionic at higher pH, takes part in the water-mediated hydrogen bond network that connects α6 and α7 from domain I with domain III (Supplementary Fig. 9b, Supplementary Table 1b). We therefore asked whether or not suppression of the pH-sensitivity of the network would stabilize the monomer at high pH, thereby reducing the pH sensitivity of the crystals. This was indeed the case, with an SP50 (pH at which 50% of crystals are dissolved) of 12.6 ± 1.0 for crystals of the E583Q mutant, compared to 11.2 ± 1.0 for WT Cry11Aa crystals (Fig. 3a), and a dissolution profile characterized by a reduced slope with no visible plateau up to pH 14. Thus, the alteration of protonation state of residues and water molecules at the intramolecular interface between domain I and domain III may be involved in the early step of Cry11Aa crystal dissolution. In Cry11Ba (G587), which displays a similar SP50 of 11.5 (Supplementary Fig. 11), this residue is substituted for glycine suggesting a different mechanism of pH-induced intramolecular separation of domain I and domain III, in Cry11Ba – or at least the involvement of additional residues at the interface.

Crystal contacts were also investigated. We first tampered with the interface enabling the honey-comb brick-wall piling of Cry11Aa tetramers (Fig. 2c, interface #2), by introducing a F17Y substitution, intended to induce electrostatic repulsion with the negatively charged D180 (distance D180(OD1) - F17(CZ) of 3.3 Å), due to deprotonation of its hydroxyl group upon pH increase (Supplementary Fig. 9c). As expected, crystals of the F17Y mutant were found to be more sensitive to increases in pH, with crystals starting to dissolve at pH as low as ~9.5 and an SP_50_ of 10.6 ± 1.0 (Fig. 3a). The dissolution profile of F17Y crystals is again characterized by a reduced slope, as compared to WT crystals, explaining that the plateau is nonetheless reached at the same pH (~pH 11.6). Nevertheless, the result suggests that dissolution of Cry11Aa crystals can be accelerated by separation of the tetramers associated through interface #2. The F17Y mutation was also found to challenge crystal formation, yielding crystals far smaller than their WT counterparts. We note that F17, D180 and the H-bond between them are strictly conserved in Cry11Ba; hence, the importance of interface 2 for crystal formation and dissolution could be extendable to crystals formed by Cry11Ba.

Next, we challenged the role of the dimerization interface (Fig. 2b interface #1). Recall that BSA at this interface, contributed by domain III from facing monomers, is 38% lower in Cry11Ba than in Cry11Aa. Furthermore, only 8 hydrogen bonds and 2 salt bridges support the interface in Cry11Ba, compared to 20 hydrogen bonds and 10 salt bridges in Cry11Aa. Y449 is positioned in the central part of the interface, and while not involved in direct H-bonding to other protein residues, supports a large H-bond network that interconnects waters and residues from facing monomers in the dimer (Supplementary Fig. 9d, Supplementary Table 1c). Hence, we investigated whether deprotonation of Y449 in the middle of the interface would significantly affect crystal dissolution by engineering of a Y449F mutation. Only a minor effect on crystal dissolution was observed (Fig. 3a), yet the mutation was detrimental to the protein stability (Fig. 3b), resulting in the growth of crystals of different size and shape (Fig. 3c).

Finally, we introduced a Y349F mutation in the β_pin_, hypothesizing that suppression of its pH-sensitive H-bond to E295(OE1) in the adjacent strand β2 would disturb the β_pin_ fold and destabilize the tetramerization interface (Fig. 2b interface #3, Supplementary Fig. 9e, Supplementary Table 1d), thereby triggering crystal dissolution. This expected effect was not observed, with crystals of the mutant displaying the same pH-induced dissolution profile as those of the WT. Nonetheless, smaller crystals were observed whose thermal stability was affected (Figure 3 and Supplementary Fig. 12), indicating that reduced stabilization of the turn preceding the β_pin_ not only impacts folding and stability of the toxin, but as well its piling into crystals – probably due to reduced tetramerization. Of note, Y349 is conserved in Cry11Ba where it H-bonds to P362(O).

Of all the single and double mutants we investigated, the Y349F mutation is that which results in the smallest crystals, closely followed by F17Y and E583Q. The Y449F mutant, however, exhibits the most noticeable change in shape compared to WT Cry11Aa. To evaluate the significance of these changes, we characterized the distribution in size of crystals of Cry11Aa-WT, Y449F, F17Y and E583Q using AFM (Fig. 3d). All three mutants had a significantly reduced volume compared to WT Cry11Aa, due to a reduced thickness of the crystals (Fig. 3d).

### Probing crystalline order of the Cry11Aa mutants by SFX

The presence of crystals does not necessarily infer that molecules are well arranged within them. We therefore used SFX to assess the level of crystalline order in crystals of the mutants that displayed modified solubilization or shape. Data were collected at the SPB/SFX beam line of the EuXFEL (Hamburg, Germany) from crystals delivered across the X-ray beam using a liquid microjet focused through a gas-dynamic virtual nozzle GDVN ^23^ (Table 2). All crystals were kept in water at pH 7 for the GDVN injection, and pulses were delivered at the MHz repetition rate (1.1 MHz) ^89,89,90^ using 10 Hz trains of 160 pulses, with a spacing of 880 ns apart. Data was collected on the AGIPD detector at its maximum rate of 3.52 kHz ^36^. With the notable exception of Y349F, crystals of all four single point mutants diffracted, yet unequal amounts of data were collected from each, and none from WT crystals, due to technical difficulties that arose during the experiment. This impeded a thorough comparison of the diffraction power of the various mutants, and prevented structure determination for the Y272F mutant. The structures of the other three mutants were determined, using the WT structure as a molecular replacement model for the phasing of diffraction data. We found that neither overall packing, tertiary structure nor interface formation is affected in the tested mutants at neutral pH (Supplementary Fig. 13). Of important note, these data demonstrate the feasibility of macromolecular nano-crystallography at MHz pulse rate using the brilliant micro-focused beam available at the SPB/SFX beamline of the EuXFEL.

**Table 2.** Data collection and refinement statistics of the Cry11Aa mutants.

|  | Cry11Aa-F17Y<br>pH 7 | Cry11Aa-Y449F<br>pH 7 | Cry11Aa-E583Q<br>pH 7 |
| --- | --- | --- | --- |
| PDB ID | 7QX7 | 7QX5 | 7QX6 |
| **Data collection** |  |  |  |
| Space group | *I* 2 2 2 | *I* 2 2 2 | *I* 2 2 2 |
| Cell dimensions (Å) | 57.72 ± 0.35 | 57.73 ± 0.24 | 57.76 ± 0.24 |
|  | 155.39 ± 1.49 | 155.55 ± 1.21 | 155.51 ± 0.98 |
|  | 171.66 ± 0.64 | 171.52 ± 0.57 | 171.51 ± 0.58 |
| Wavelength (Å) | 1.33 | 1.33 | 1.33 |
| X-ray beam focus (μm) | 1.3 | 1.3 | 1.3 |
| No. collected frames | 3150500 | 5993679 | 3523741 |
| No. indexed frames | 28227 | 104359 | 21833 |
| No. merged crystals | 28811 | 111014 | 22760 |
| Resolution range (Å) | 23.17 – 3.40 | 23.78 – 3.10 | 23.50 – 3.30 |
|  | (3.40 – 3.48) | (3.10 – 3.17) | (3.30 – 3.38) |
| No. observations | 2908715 (141787) | 20279640 (1092683) | 3210163 (154933) |
| No. unique reflections | 10990 (707) | 14447 (950) | 12014 (787) |
| <I/σ(I)> | 6.31 (1.67) | 9.95 (1.35) | 5.64 (1.52) |
| R<sub>split</sub> (%) | 19.74 (76.86) | 11.79 (89.56) | 21.11 (80.18) |
| CC<sub>1/2</sub> | 0.96 (0.21) | 1.00 (0.60) | 0.99 (0.31) |
| Completeness (%) | 99.6 (100.0) | 99.7 (100.0) | 99.6 (100.0) |
| Multiplicity | 265.7 (200.5) | 1403.7 (1150.2) | 267.2 (196.8) |
| **Refinement** |  |  |  |
| Resolution range (Å) | 23.17 – 3.40 | 23.18 – 3.10 | 23.08 – 3.30 |
| No. reflections | 10986 | 14442 | 12008 |
| R<sub>work</sub>/R<sub>free</sub>* | 21.2 / 25.1 | 22.4 / 25.2 | 21.5 / 25.4 |
| No. atoms |  |  |  |
| Protein | 4970 | 4965 | 4970 |
| Water | 5 | 13 | 6 |
| B-factors (Å<sup>2</sup>) |  |  |  |
| Main chain | 54.6 | 43.1 | 45.4 |
| Side chain | 54.2 | 42.7 | 45.3 |
| Water | 52.9 | 59.3 | 36.0 |
| R.m.s.d. |  |  |  |
| Bonds lengths (Å) | 0.002 | 0.002 | 0.003 |
| Bonds angles (°) | 0.448 | 0.441 | 0.489 |

\* R<sub>free</sub> is calculated using 5% of random reflections excluded from refinement.

The needle shape inclusions formed by C11AB were also investigated by SFX and found to present some crystalline order, as evidenced by diffraction rings up to ~6 Å resolution in the powder diagram calculated from the maximum projection of 395656 hits (Supplementary Fig. 8d). It is clear, however, that a high-resolution structure is not readily practicable with these crystals, either because their small size makes them unsuitable for diffraction using a micro-focused XFEL beam or due to intrinsic disorder.

## Discussion

We here report the previously-unknown structures of Cry11Aa and Cry11Ba, the two most potent Cry δ-endotoxins expressed by mosquitocidal *Bti* and *Btj*, respectively. Both toxins occur as natural nanocrystals that are produced during the sporulation phase of the bacteria, and dissolve upon elevation of pH in the mosquito larvae gut. Proteolytic activation enables binding to their specific receptors ^37^, including a membrane embedded alkaline phosphatase^38^ but as well the co-delivered Cyt1Aa ^12,39–41^, triggering insertion in gut cell membranes and subsequent oligomerization into pores that will eventually kill the cells. Both toxins are of industrial interest due to their environmental safety, explained by the multi-step activation outlined above, and to their high stability as crystals. Our results shed light on the mechanisms of *in vivo* crystallization, pH-induced dissolution and proteolytic activation, and on structural features that support the toxins specificity with respect to other Cry toxins. Thereby, our work offers a foundation for further improvement of the toxic activity or crystal size by rational design. Additionally, we demonstrate the feasibility of *de novo* structure determination of a previously-unknown protein-structure by SFX, from nanocrystals only 10,000 unit-cells across, using a single caged-terbium (TbXo4) derivative. Below, we recapitulate these findings and discuss their implications.

### In vivo *crystallization pathway of Cry11 toxins*

The building block of Cry11Aa and Cry11Ba crystals is a tetramer formed by the interaction of two dimers, via their domain II. The dimers are themselves assembled from the interaction of two monomers, via their domains II and III. Crystals form from the honey-comb brick-wall piling of tetramers, as enabled by the face-to-back interaction of domain I from symmetry-related tetramers (Figure 2). Thus, all three domains are involved in the *in vivo* crystal packing of Cry11 toxins, each contributing a two-fold axis. This observation contrasts with other toxin structures determined from *in vivo* grown crystals, wherein either propeptide(s) (e.g. *Lysinibacillus sphaericus* BinAB ^28^ and *Bti* Cyt1A ^12^) or a specific domain (e.g. domain I in *Btt* Cry3Aa from ^42,43^) serves as the major contributor to crystallization. Expanding to previously determined Cry δ-endotoxins ^12,28,42,44^ structures, solved from *in vitro* grown macrocrystals obtained following dissolution of the natural crystals at high pH, the same trend is observed – *i.e*., crystallization mostly depends on a dedicated portion of the protein, either it be a N-terminal and/or C-terminal propeptide (e.g., the ~650 C-terminal residues in *Btk* Cry1Ac) or a specific domain (e.g. domain II in *Btk* Cry2Aa). Thus, the Cry11Aa and Cry11Ba structures illustrate a yet unobserved pathway for *in vivo* crystallization, wherein all domains act on a specific step of the coalescence process, viz. dimerization (domains II and III from two Cry11 monomers), tetramerization (domains II from two Cry11 dimers) and tetramer-piling (domains I in each tetramer). With Cry11Aa featuring a larger dimerization interface, and Cry11Ba a larger interface between piled tetramers, the two structures underline different levels of tradeoff between packing *into* tetramers and packing *of* the tetramers.

The difference in thickness of Cry11Aa and Cry11Ba crystals is of interest. Considering that all crystals were produced in *Bti*, we could exclude the possibility that the slightly larger size of Cry11Ba crystals originates from a more efficient crystallization machinery in *Btj* than *Bti*. Puzzled by the presence of a 77-residue long low complexity region at the C-terminus of Cry11Ba (LCR-Cry11Ba), which is absent in Cry11Aa, we asked whether or not a C-terminal fusion of LCR-Cry11Ba with Cry11Aa would result in larger crystals. LCR regions have indeed been shown to support a variety of functions, including chaperoning ^35^ and reversible oligomerization ^45,46^ so that a role in crystal nucleation and/or growth could not be excluded. Support of the first, but not the second hypothesis was obtained. Indeed, the C11AB chimera, consisting of a fusion of LCR-Cry11Ba to the C-terminus of Cry11Aa, yields smaller crystals that poorly diffract, even when exposed to high intensity XFEL pulses. This observation is in line with previous results which showed that substitution of Cry11Ba domain III for that of Cry11Aa leads to limited expression and comparatively small inclusions ^47^. Thus, the LCR region of Cry11Ba is unlikely to account for the difference in size between Cry11Aa and Cry11Ba crystals. Instead, we favor the hypothesis that it is the larger surface of interaction between piled tetramers that accounts for the larger size of the Cry11Ba crystals. Given the absence of electron density for LCR-Cry11Ba residues in the Cry11Ba structure, and the abundance of needle-like inclusions in the parasporal body enveloping the C11AB crystals, it is reasonable to assume that they do not engage in structurally important interactions with functional domains, but rather favor nucleation of crystals. This aid-to-nucleation would be required for Cry11Ba, which features a reduced dimerization interface, but not for Cry11Aa, wherein this interface is 62 % larger. In line with this hypothesis, four regions are predicted to form short adhesive motifs of the Low Complexity, Amyloid-like Reversible Kinked Segments (LARKS) type (Supplementary Fig. 3).

### Cry11 toxins depart from the canonical Cry δ-endotoxins architecture

The structures of Cry11Aa and Cry11Ba shed light on features that would not have been predicted based on sequence alignments (*i.e*., by homology modelling), and which largely deviate from the canonical organization observed in other Cry δ-endotoxins ^12,28,42,44^. The most notable difference is the presence of a ~36 to 38 residue insertion between strands β4 and β5 in domain II, which results in an extra β-strand, coined β_pin_. The β_pin_ not only participates in the formation of a modified β-prism, but contributes to a two-fold axis that supports tetramerization of Cry11 toxins through formation of two large β_h_-β4-β3-β2-β_pin_ – β_pin_-β2-β3-β4-β_h_ sheets between symmetry-related dimers into a tetramer. The observation of proteolytic cleavage sites at both the N- and C-termini of the β_pin_ suggests that it is removed upon activation by mosquito gut proteases, in line with the observation of ~32 and ~36 kDa fragments upon proteolytic activation of the Cry11 toxins ^32^. If true, the unique role of the β_pin_ would be to support *in vivo* crystallization and its removal would entail the dissociation of tetramers into dimers and eventually monomers. While mutagenesis results indicate that this interface does not play a major role in crystal dissolution (see below), it seems likely that upon pH elevation and deprotonation of tyrosines and acidic groups, electrostatic repulsion will occur between Y349(OH) and E295(OE2) in Cry11Aa, and between Y350(OH) and P362(O) in Cry11Ba. Increased disorder of these regions could facilitate the access of proteases, and thus favor the activation of the Cry11Aa and Cry11Ba toxins. This hypothesis would rationalize the reluctance of the two toxins to recrystallize *in vitro* after pH induced dissolution, due to an impossibility to re-form tetramers – or at least, to re-match the exact positioning of the β-pin. The Cry11 toxins also differ from other Cry δ-endotoxins by the presence of a conserved N/D-DDLGITT insertion between β21 and β22, contributing a short helix, and by deletions of ~5-10 residues in the α3-α4 and β20-β21 loops, respectively. Compilation of these changes likely explains failures to phase the Cry11 structures by the molecular replacement method, even when *Btk* Cry2Aa, which also features a α_h_β_h_-handle, was used as a starting model.

### Mapping the interfaces involved in crystal dissolution

Our efforts to determine the structures of Cry11Aa and Cry11Ba at alkaline pH were unsuccessful, due to high sensitivity of crystals diffraction quality to pH increase. In the case of Cry11Aa we could not collect data, while in the case of Cry11Ba, we obtained a low-resolution structure which, while showing possible inter-domain rearrangements, did not inform on specific side chain rearrangements. Therefore, we resorted to site-specific mutagenesis to obtain information regarding the crystal dissolution pathway. We found that the crystal interface most sensitive to pH elevation is the one enabling the honey-comb brick-wall piling of Cry11 tetramers, with the Cry11Aa-F17Y mutant displaying increased pH sensitivity (with an SP_50_ of 10.6 ± 1.0 compared to 11.2 ± 1.0 for WT Cry11Aa crystals). In contrast, the dimerization (Y349F mutant) and tetramerization interfaces (Y449F mutant) appear to be less pH-sensitive. At the monomer level, we found that the three-domain interface to which α_8_ and the α_h_β_h_-handle contribute is not very sensitive to pH increase (Y272Q and D507N+D514N mutants), possibly due to burying of mutated residues at the interface, preventing bulk solvent to access these sites. Alternatively, interaction of Cry11 toxins with its membrane-bound receptors ^37^ could be a required step to expose α_8_, shown to play a major role in binding and toxicity ^48^.

The intramolecular domain I vs. domain III interface was found to be important for the pH-induced crystal dissolution, with the Cry11A E583Q mutant displaying a reduced sensitivity to pH (SP_50_ of 12.6 ± 1.0). Yet unlike the other tested interfaces, which are overall well conserved, the domain I vs. domain III interface differs in Cry11Aa and Cry11Ba, suggesting that caution is advised upon reflecting results obtained from Cry11Aa mutants onto Cry11Ba. Indeed, E583 is substituted for glycine in Cry11Ba (G587), suggesting a different mechanism of pH-induced separation of domain I and domain III – or at least, the participation of other residues. Structural analysis suggests that the substitution of Cry11Ba Q247 for a glutamic acid could compensate for the absence of E583, enabling electrostatic repulsion of V494 (β14) – found at the opposed end of this interface – upon pH elevation. Numerous other residues at this interface, otherwise mostly conserved between Cry11Aa and Cry11Ba, remain as candidates to further tune the pH sensitivity. For example, Y241(OH) is H-bonded to D586(OD1; 2.6 Å) and D590(OD2; 2.8 Å) in Cry11Aa and Cry11Ba, respectively, suggesting that mutation of this residue into a phenylalanine (Y241F) and/or of D586/D590 into asparagines would reduce the pH sensitivity while not affecting stability. Likewise, E234 H-bonds to Q625(NE2; 2.6 Å) in Cry11Aa, and to K629(NZ; 2.8 Å) and R553(NH1; 2.9 Å) in Cry11Ba, suggesting that a E234Q mutation would reduce pH sensitivity in the two toxins whilst not affecting their folding. Inversely, the mutation into a glutamic acid of Q511/Q515, squeezed between a tryptophan (W584/W588), an arginine (R549/R553) and a glutamic acid (E234), would be expected to increase the pH sensitivity of the domain I vs. domain III intramolecular interface in both Cry11Aa/Cry11Ba – and by extension, that of their crystals.

### Implication for the future of nanocrystallography using SFX

In this study, *de novo* phasing was required – not because of the absence of homologous structures, but because none of those available were sufficiently close to serve as a search model for molecular replacement. Using Tb-Xo4, a caged terbium compound, we could phase the *Bti* Cry11Aa structure by SAD, from ~77,000 diffraction patterns collected on crystals only 10,000 unit cell across – an achievement to compare to the determination of the structure Ls BinAB from > 370,000 patterns (native and three derivatives) collected on crystals 100,000 unit cell across ^49^. Our success in phasing the Cry11Aa structure stemmed from a combination of advances in SFX data processing tools over the last five years and the use of a dramatically powerful phasing agent, and should offer hope to investigators seeking to determine the structure of proteins of which no known structural homologue exists and that have to resort to SFX due to smallness of their crystals. It is foreseeable, however, that *de-novo* structure determination will be helped by recent advances in comparative and *ab-initio* modelling and the availability of programs such as RosettaFold ^50^ and AlphaFold2 ^51^, capable of producing a decently-accurate structure for virtually all proteins and thus a good model for phasing of crystallographic data by molecular replacement. Latest releases of the two programs were published in the final stage of the writing of this manuscript, hence we asked whether or not the availability of these tools would have facilitated our journey towards the Cry11 toxins structures, and submitted the sequence of Cry11Aa to the two servers. For RosettaFold, the rmsd to the final refined structure of the five best models was over 4 Å, with discrepancies observed mostly in domain II. For AlphaFold2, however, the two first models displayed rmsd of 1.2 and 1.0 Å to the final structure, respectively. Using the worst of these two models, we could find a molecular replacement solution using Phaser, and a partial model featuring 95% of the residues in sequence was obtained after 20 cycles of automatic iterative model-building and refinement using Bucanneer ^52^ and Refmac ^53^. Thus, a problem which occupied a handful of crystallographers for several years could have been solved in less than an hour using the new tools recently made available to the structural biology community. Based on our results, it is tantalizing to claim that the phase problem in crystallography has been solved, or that experimental structural biology has lived, but such assertions would likely be shortsighted. Rather, we encourage investigators to challenge AlphaFold2 and RosettaFold as much as humanly possible, but to not forsake *de novo* phasing as it may remain the only route to success in difficult cases where molecular replacement based on such models does not work ^54^. It must also be emphasized that in the case of Cry11 toxins and, more generally, naturally-crystalline proteins, the issue is not just phasing, but packing. For such proteins, crystal formation and dissolution serve function, hence characterization of packing interfaces is central to finely comprehend their bioactivation cascades. Without the naturally-occurring crystals and the atomic resolution SFX structures, it would not have been possible to make predictions on potential mutations affecting Cry11Aa crystal formation or dissolution.

## Materials and Methods

### Crystal production and purification

Crystals of Cry11Aa and Cry11Ba were produced by electrotransformation of the plasmids pWF53 and pPFT11S ^55^ into the acrystalliferous strain 4Q7 of *Bacillus thuringiensis* subsp. *israelensis* (Bti; The Bacillus Genetic Stock Center (BGSC), Columbus OH, USA), respectively ^56^. Colonies were selected on LB agar medium supplemented with erythromycin (25 μg/mL) and used to inoculate precultures of LB liquid medium. For Cry11Aa production, precultures were spread on T3 sporulation medium. After incubation at 30°C for 4 days, spores/crystals suspensions were collected using cell scrapers and resuspended in ultrapure water. After sonication-induced cell lysis and subsequent centrifugation at 4000 g for 45 min to discard cell and medium debris, pellets were resuspended in water and crystals were purified using a discontinuous sucrose gradient (67-72-79 %). After ultracentrifugation, crystals were recovered and several rounds of centrifugation/resuspension in ultrapure water allowed discarding as much sucrose as possible for proper downstream application. Crystal purity was verified by SDS-PAGE on 12% gels. Purified crystals were conserved in ultrapure water at 4 °C until use. For Cry11Ba, a glycerol stock of the 4Q7/pPFT11S was streaked onto 25 μg/mL erythromycin Nutrient Agar plates. From here a single colony was selected and added to a Glucose-Yeast-Salts (GYS) media culture and allowed to grow continuously at 30°C, 250 rpm for 5 days. This culture was then spun down, resuspended in ultrapure water, and the lysate was sonicated for 3 min at 50% duty. The sonicated lysate was added to the 30-65% discontinuous sucrose gradient (35-40-45-50-55-60-65 %) and spun down for 70 min at 20,000 rpm and 4°C. The sucrose gradient was then hand fractionated with Cry11Ba crystals collected around 57-60% and dialysed into ultrapure water. Crystal characterization and purity was completed by phase contrast light microscopy, X-ray powder diffraction, transmission electron microscopy, and 4-12% SDS-PAGE gels. The pure Cry11Ba crystals were stored at 4°C in ultrapure water.

### Cry11Aa mutagenesis

Based on the SFX structure of Cry11Aa, a total of 7 mutants of Cry11Aa were constructed to challenge the different crystal packing and intramolecular interfaces. The rationale behind these mutations is illustrated in Supplementary Fig. 9 and discussed in the main text. Point-mutations were inserted into *cry11aa* gene by Gibson assembly using pWF53 as a backbone ^56^. Two different primer couples were used for each mutation to amplify two fragments that were complementary by their 15-18 bp overlapping 5’ and 3’ overhangs with a target Tm of 50°C. Point mutations were inserted in the complementary part of the overhangs of the two fragments spanning the *cyt1aa* region to be mutated. The double mutant D507N-D514N was successfully constructed in a single-step by respectively adding the D507N mutation on the non-overlapping overhang region of the forward primer, and the D514N on the non-overlapping overhang of the reverse one. The triple mutant Y272Q-D507N-D514N was constructed by using the primers containing the Y272Q mutation and the plasmid pWF53-D507N-D514N as a backbone. In addition to the point mutants, a Cry11Aa-Cry11Ba chimeric toxin – coined C11AB – was also constructed. For this, the sequence of the *cry11aa* gene was fused with the 234 bp extra 3’ extension of *cry11ba* gene, which is suggested to feature a low complexity region (LCR) based on sequence analysis using the LCR-eXXXplorer web platform (http://repeat.biol.ucy.ac.cy/fgb2/gbrowse/swissprot) ^57^, which implements the CAST ^19^ and SEG ^20^ computational methods to identify LCR. The C11AB chimera was constructed by Gibson assembly following a “1 vector, 2 fragments” approach. The plasmid pWF53 containing the *cry11aa* gene was used as a backbone and the *cry11ba* 3’ fragment was amplified from the extracted and purified plasmid of the WT strain of *Btj* containing the *cry11ba* gene. The list of primers used for plasmids construction is available in Supplementary Table 2. For each plasmid construction, the fragments with overlapping overhangs were assembled using the NEBuilder HiFi DNA Assembly (New England BioLabs) as previously described ^12^. Briefly, after 90 min incubation at 50°C, the constructed plasmids were transformed by heat shock into chemically competent Top10 *Escherichia coli* (New England BioLabs). Plasmids were extracted from colonies selected on LB agar medium containing ampicillin (100 μg mL^-1^) using the NucleoSpin Plasmid extraction kit (Macherey-Nagel) following the manufacturer’s instructions. The successful construction of each plasmid was assessed by double digestion (EcoRI and BamHI) followed by migration on 1% agarose gel stained with SYBR Safe (Invitrogen) and by Sanger sequencing of the region containing the mutation at the Eurofins Genomics sequencing platform. Of note, the *cry11aa* gene was also fully sequenced to validate its sequence for mutagenesis primer design and for comparing the expected toxin size to the observed one in mass spectrometry analyses. All mutants were produced as crystals in *Bt*, as described above. The presence of the mutated *cry11aa* gene sequence in the transformed *Bt* colony used for production was verified by colony PCR using specific primers and Sanger sequencing at the Eurofins Genomics sequencing platform. Crystals from all mutants were analyzed by SDS-PAGE on 12% gels. For C11AB, its proper size was confirmed by using the “gel analysis” module implemented in the software ImageJ v1.51k (*N* = 7) ^58^.

### Crystal visualization by scanning electron microscopy (SEM)

Purified crystals of Cry11Aa WT and of the 7 mutants were visualized using either a Zeiss LEO 1530 scanning electron microscope from the SEM facility of the European Synchrotron Radiation Facility (ESRF, Grenoble, France), a Thermo Fisher Quanta 650 FEG environmental SEM (ESEM) available for users at the European XFEL (EuXFEL, Hamburg, Germany) or a JEOL JSM-6700M FE-SEM (UCLA, Los Angeles, USA). For SEM at ESRF, samples were coated with a 2 nm thick gold layer with the Leica EM ACE600 sputter coater before imaging. For ESEM at the European XFEL, samples were diluted (1/1000) and mixed with 25 mM of ammonium acetate. Samples were then coated with a thin gold layer as described above using a Leica EM ACE600 sputter coater as well. Images were recorded at 15 kV acceleration voltage by collecting secondary electrons using an Everhart-Thornley-Detector (ETD detector) in high-vacuum mode. For SEM at UCLA, samples were diluted (1/5) and ultrapure H2O.they were then added to 300 mesh Cu F/C grids that were positively glow discharged. These samples were then wicked away and washed with ultrapure water, wicked, and allowed to dry overnight to ensure all moisture had evaporated inside of a dessicator. These were then attached to a holder with carbon tape and coated with an Anatech Hummer VI sputter coater with approximately 2 nm of thick gold layer. Images were recorded at 5 kV acceleration voltage by collecting secondary electrons using a Lower secondary electron (LEI) or Upper secondary electron in-lens (SEI) detector.

### Crystal visualization by transmission electron microscopy (TEM)

Non-purified crystals of Cry11Aa WT were visualized using a Thermofisher TF20 electron microscope from the IBS electron microscopy platform. For negative staining TEM, samples were diluted 5 times in H_2_O and 4 μl of the diluted sample was introduced to the interface of an amorphous carbon film evaporated on a mica sheet. The carbon film was then floated off the mica sheet in ~200 μl 2% sodium silicotungstate (SST) solution. The carbon film with the crystal sample was then recovered onto a Cu 300 mesh TEM grid after 30 s, let dry, and imaged at 200 keV. Images were recorded on a Gatan OneView CMOS detector. Non-purified crystals of Cry11Ba WT were visualized using an FEI Tecnai T12 electron microscope within the UCLA California Nanoscience Institute, EICN facility. For negative staining TEM, samples were prepared by adding 5 μL of pure crystal fractions in 10 μL ultrapure H2O. 2.5 μL of this sample was added to 300 mesh Cu F/C grids that were positively glow discharged. These samples were then wicked away using Whatman 1 filter paper; washed with 2.5 μL ultrapure H2O, wicked; and negatively stained with 2.5 μL 2% uranyl acetate, wicked. These were allowed to dry overnight to ensure all moisture had evaporated and imaged at 120 keV. Images were recorded on a Gatan 2kX2k CCD.

### Crystal characterization by atomic force microscopy (AFM)

Crystals of Cry11Aa were visualized by AFM as previously described^12^. Briefly, 5 μL of crystals suspended in ultrapure water were deposited on freshly cleaved mica. After 30 min in a desiccation cabinet (Superdry cabinet, 4% relative humidity), crystals were imaged on a Multimode 8, Nanoscope V (Bruker) controlled by the NanoScope software (Bruker, Santa Barbara, CA). Imaging was done in the tapping mode (TAP) with a target amplitude of 500 mV (about 12 nm oscillation) and a variable setpoint around 70% amplitude attenuation. TESPA-V2 cantilevers (k = 42 Nm^-1^, Fq = 320 kHz, nominal tip radius = 7 nm, Bruker probes, Camarillo, CA, USA) were used and images were collected at ~1 Hz rate, with 512- or 1024-pixel sampling. Images were processed with Gwyddion ^59^, and if needed stripe noise was removed using DeStripe ^60^. Measurements were performed on Cry11Aa WT and on mutants selected on the basis of their aspect in eSEM images (Y449F) or their solubilization pattern (F17Y and E583Q). Size measurements were performed on AFM images using Gwyddion ^59^ in a semi-automated protocol. A classical height threshold was applied to each image to select as many individual crystals as possible. Sometimes, partially overlapping crystals were individualized using the manual edition of the mask of selected crystals by adding a separation line. Finally, a filter was applied to remove very small selections (artefacts) or crystals touching the edge of the image. Measures were obtained using the ‘distribution of grains’ feature in Gwyddion where the crystal thickness (T) is the returned mean value, the volume (V) is the Laplacian background basis volume, and the length and width are the major and minor semi-axes of equivalent ellipses, respectively. The total number of crystals measured are: 45 for WT, 93 for F17Y, 60 for Y449F, and 94 for E583Q.

### Data collection history

The Cry11Aa/Cry11Ba structure determination project was initiated in 2015. Data were collected at five different occasions, in two XFEL facilities, namely at the Linac Coherent Light Source (LCLS), Stanford (USA) and EuXFEL, Hamburg (Germany). During our first LCLS-SC3 beamtime (cxi04616), we collected data from native Cry11Ba (2.3 Å resolution), and in our second (LO91), we collected data from native Cry11Aa (2.8 Å resolution). Nanocrystals grown by recombinant expression in the modified acrystalliferous 4Q7 strain of *Bti* were injected by a microfluidic electrokinetic sample holder (MESH) device^61^ in the microfocus chamber of LCLS-SC3 ^62^. Crystals were. After data reduction using cctbx.xfel and dials (hit-finding through merging) ^63–66^, we attempted phasing of both datasets by molecular replacement (MR), using sequence-alignment based multi-model approaches implemented in Mr Bump (based on MR by Molrep ^67^) as well as custom-scripts testing models produced by Rosetta ^68^ (using the Robetta server; http://robetta.bakerlab.org/) and SwissProt ^25^ (https://www.ebi.ac.uk/uniprot/) servers (based on MR by Phaser ^69^). Failure to find a homolog of a sufficiently-close structure led us to attempt de novo phasing of the Cry11 nanocrystalline proteins. Initially, we aimed at obtaining experimental phases for Cry11Ba, considering that its larger crystals would produce a stronger diffraction signal which in turn would facilitate phasing. Hence, we collected derivative data on Cry11Ba, from crystals soaked with Gd, Pt and Au salts (P127 experiment) before injection using a MESH device^61^. Unfortunately, the data did not allow phase determination, as indicated by very weak and absent peaks in the isomorphous and anomalous difference maps, respectively (Supplementary Fig. 2), due to low occupancy of the soaked metal ions. Hence, we shifted focus to Cry11Aa crystals soaked with a recently introduced caged-terbium compound, Tb-Xo4 ^16^ (P125 experiment). Crystals were injected using a GDVN ^23^ liquid microjet in the microfocus chamber of LCLS-SC3 ^62^. Online data processing was performed using NanoPeakCell ^70^ and CASS ^71^. Offline data processing with NanoPeakCell ^70^ (hit finding) and CrystFEL ^72^ (indexing and merging) revealed a strong anomalous signal that enabled determination of the substructure and phasing of the SFX data, using Crank2 ^73^ and its dependencies in the CCP4 suite ^74^ (see below for more details). The Cry11Aa structure was thereafter used to phase the Cry11Ba datasets by molecular replacement, revealing *a posteriori* that the Gd, Pt and Au ions had successfully bound to the crystalline Cry11Ba in the various derivatives collected during P127, despite anomalous and isomorphous signals being too weak to enable phasing. We last attempted data collection on Cry11Aa and Cry11Ba crystals soaked at elevated pH and injected by a MESH device (P141 experiment). Only Cry11Ba crystals could sustain the pH jump and yielded usable data. From the comparative analysis of the Cry11Aa and Cry11Ba structures, we nonetheless designed mutations aimed at increasing or decreasing the resilience of crystals; these were introduced in the Cry11Aa gene, and crystals were produced by recombinant expression in *Bti*. From these, SFX data were collected at the MHz pulse rate, during experiment P2545 at the SPB/SFX beam line of EuXFEL where a GDVN was used to inject crystals. The data were also processed with NanoPeakCell ^70^ (hit finding) and CrystFEL ^72^ (indexing and merging).

### Data collection and processing, and structure refinement

During the P125 beamtime at LCLS, where the SAD data used for the phasing of the Cry11Aa structure were collected, the X-ray beam was tuned to an energy of 9800 eV (i.e. a wavelength of 1.27 Å), a pulse duration of 50 fs, a repetition rate of 120 Hz, and a focal size of 5 μm. SAD data were collected from nanocrystals soaked for 30 hours with Tb-Xo4 at 10 mM in water, prior to GDVN injection ^23^. Of 558747 images collected using the 5 μm beam available at the at LCLS-SCC, a total of 77,373 images were indexed of which 76,687, 292, 217 and 177 using Xgandalf ^75^, Dirax ^76^, taketwo ^77^ and Mosflm ^78^, respectively, in CrystFEL v.0.8.0 ^79^. Post-refinement was not attempted, but images were scaled one to another using the ‘unity’ model in CrystFEL *partialator*, yielding a derivative dataset extending to 2.55 Å resolution. *A posteriori*, we found that simple Monte Carlo averaging using the ‘second-pass’ option in CrystFEL *process_hkl* would have yielded data of similar quality. A native dataset was also collected and processed in the same fashion yielding, from 792623 collected patterns of which 48652 were indexed, a dataset extending to 2.60 Å resolution. The substructure of the derivative dataset was easily determined by ShelxD (figure of merit (FOM): 0.22), prompting us to try automatic methods for structure determination. Using Crank2 ^73^ and its dependencies (ShelxC, ShelxD, Solomon, Bucanneer, Refmac5, Parrot) in CCP4 Online ^80^, the FOM was 0.52 after density modification, and rose to 0.88 upon building of 613 residues. This first model was characterized by R_work_/R_free_ of 27.7/32.1 % and was further improved by automatic and manual model building in phenix.autobuild ^81^ and Coot ^82^ until 630 residues were correctly build. This model was then used to phase the native data. Final manual rebuilding (using Coot ^82^) and refinement (using phenix.refine ^83^ and Refmac5 ^53^) afforded a native model characterized by R_work_/R_free_ of 17.2/24.1 % and consisting of most of the 643 residues. Only the first 12 N-terminal residues are missing (Table 1).

Cry11Ba data were collected during the cxi04616 and P141 beamtimes at LCLS-CXI. At both occasions, the photon energy was 9503 eV (i.e., a wavelength of 1.30 Å), a pulse duration of 50 fs, a repetition rate of 120 Hz, and a focal size of 1 μm – *i.e*., a similar standard configuration (pulse length, repetition rate) than that used for Cry11Aa, notwithstanding the beam size and wavelength. Data were collected from crystals at pH 6.5 (30% glycerol in pure water; cxi04616) and pH 10.4 (30% glycerol in 100 mM CAPS buffer; P141), presented to the X-ray beam using a MESH injector ^22^. Of 813133 images collected for the pH 6.5 dataset, 19708 were indexed and scaled, post-refined, and merged using cctbx.xfel^63–66^ and PRIME^84^, yielding a dataset extending to 2.2 Å resolution. The Cry11Aa structure was used as a starting model to phase the Cry11Ba pH 6.5 dataset by molecular replacement using Phaser ^69^ with initial R_work_/R_free_ being 34.4/40.4 %. Manual model building (using Coot ^82^) and refinement (using Refmac ^53^ and Buster ^85^) afforded a model characterized by R_work_/R_free_ of 18.9/23.8 % (Table 1). Because the 2.55 Å resolution Cry11Ba pH 10.4 data was of limited utility, in view of absence of major peaks in the Fourier difference map calculated with the pH 6.5 data as a reference, and of a 1% change in the unit cell volume ascribable to the use of a different glycerol concentrations during injection of the two samples, it was not included in our PDB and CXIDB depositions.

Diffraction data on the Cry11Aa mutants at pH 7.0 was acquired on the SPB/SFX beamline at EuxFEL during our P002545 beamtime allocation, using a GVDN injector and X-ray energy and focal size of 9300 eV (1.33 Å) and 1.3 μm (FWHM), respectively. Technical problems allowed us to collect only a limited number of diffraction pattern of the Cry11Aa-Y349F mutant. 3,150,500; 5,993,679 and 3,523,741 images were collected for the F17Y, Y449F and E583Q mutant, respectively, of which 28,227; 104,359 and 21,833 could be processed using CrysFEL0.8.0 ^79^ and MonteCarlo based scaling and merging. The three structures were solved using MR with Phaser ^69^, using the Cry11Aa WT structure as input model. The structures were refined using Phenix.refine ^83^ and Coot ^82^, with final R_work_/R_free_ values of 21.2/25.1 % for Cry11Aa-F17Y, 22.4/25.1 % for Cry11Aa-Y449F and 21.5/25.4 % for Cry11Aa-E583Q (Table 2).

### Structure analysis

Figures were prepared using pymol v. 2.5^86^ (Fig. 1, 2 and Supplementary Fig. 4, 9, 13) and aline (Supplementary Fig. 3) ^87^. Radii of gyration were predicted using the pymol script rgyrate (https://pymolwiki.org/index.php/Radius_of_gyration). Interfaces were analyzed with PISA ^88^ and rmsd among structures were calculated using pymol using the ‘super’ algorithm. Sequence based alignment – performed using EBI laglign and ClustalW ^89^ – was challenged by the large gaps between Bti Cry11Aa, Btj Cry11Ba, Btk Cry2AA and Btt Cry3Aa, while structure-based alignment – performed using SSM ^90^ – was blurred by the varying size of secondary structure elements in the three domains of the various toxins. Hence, for Supplementary Fig. 1, 3, the alignment of Bti Cry11Aa, Cry11Ba, Cry2AA and Cry3Aa was performed using strap ^91^ which takes into account both sequence and structural information. Specifically, the online version of the program was used (http://www.bioinformatics.org/strap/) ^92^.

### Structure prediction using AlphaFold2 and RosettaFold

RosettaFold ^50^ predictions were obtained by submitting the sequence to the Rosetta structure-prediction server (https://robetta.bakerlab.org). AlphaFold2 ^51^ predictions were obtained by use of the Colaboratory service from Google Research (https://colab.research.google.com/github/sokrypton/ColabFold/blob/main/beta/AlphaFold2_advanced.ipynb). The mmseq2 method ^93,94^ was employed for the multiple-sequence alignement instead of the slower jackhmmer method ^95,96^ used in ^51^. Structural alignments were performed using the *align* tool in PyMOL ^86^. Molecular replacements trials were carried out with Phaser ^69^. Using the best five RosettaFold models, all characterized by an overall rmsd to the final structure superior to 4 Å, no molecular replacement solution could be found, due to inaccurate prediction of domain II β_pin_ and α_h_-β_h_ regions, resulting in clashes. The best alphafold2 model was yet successful at predicting the domain II structure, which enabled successful phasing by molecular replacement, yielding a model characterized by R_free_ and R_work_ values of 0.322 and 0.292, respectively. This model was further use as a starting model for automatic model building and refinement using the buccaneer pipeline in CCP4, resulting in a model characterized by R_free_ and R_work_ values of 0.245 and 0.215, respectively, after only five automatic cycles of iterative model-building, refinement and density modification using bucanneer ^52^ and refmac5 ^53^ in the CCP4 suite ^74^.

### Crystal solubilization assays

The solubility of crystals of Cry11Aa WT and of the mutants F17Y, Y272Q, Y349F, Y449F, D507N-D514N and E583Q was measured at different pH values as previously described ^12^. Briefly, crystal suspensions were centrifuged at 11,000 g for 2 min and resuspended in 18 different buffers with pH ranging from 8.6 to 14.2. After 1h incubation in each buffer, crystals were centrifuged and the supernatant was collected. The concentration of soluble toxin in the supernatant was quantified using a Nanodrop 2000 (Thermo Fisher Scientist) by measuring the OD at 280 nm and by using the molar extinction coefficient and toxin size (102,930 M^-1^ cm^-1^ and 72.349 kDa, respectively, as calculated with the ProtParam tool of ExPASy (https://www.expasy.org) using the Cry11Aa protein sequence available under accession number “P21256 [https://www.uniprot.org/uniprot/ P21256]”). Solubility was measured in triplicate for each toxin (Cry11Aa WT and mutants) and each pH. Data are normalized and represented as percentage of solubilization by dividing the concentration measured at a given pH by the concentration at the highest pH measured. Solubility of Cry11Aa WT and its different mutants was compared by calculating SP50 (pH leading to solubilization of 50% of crystals) as previously described ^12^, by fitting the data using a logistic regression model for binomial distribution using a script modified from ^97^. Differences in SP50 between mutants were considered significant when 95% confidence intervals (CI), calculated using a Pearson’s chi square goodness-of-fit test, did not overlap ^98^. All statistics were conducted using the software R 3.5.2 ^99^.

For the Cry11Ba, the crystal suspensions were centrifuged at 13,300 g for 3 min and ultrapure H2O was removed from crystals. They were then resuspended in one of 18 buffers ranging from pH 7 to 14. These crystals were incubated for 1 hr, afterwards the solution was centrifuged at 13,300 g and the supernatant was separated from the crystal pellet. The concentration of the supernatant was then quantified by a ThermoFisher Nanodrop One (Thermo) by measuring the OD for 280 nm and utilizing the molar extinction coefficient and toxin size (114600 M^-1^.cm^-1^ and 81344.18 Da respectively) that were calculated with Expasy ProtParam using the Cry11Ba sequence available at Uniprot.org under accession number Q45730. Solubility was measured in triplicate for the toxin at each pH measured. This was then further tested by conducting a turbidity assay by resuspending the crystal pellet in 150 μL ultrapure H2O and placed in a 96-well plate to be read on an NEPHELOstar Plus (BMG Labtech) nephelometer. These counts were normalized by subtracting the background signal and conducted in triplicate.

### Proteomic characterization

For SDS-PAGE experiments, samples heated to 95 °C were migrated on 12 % SDS-PAGE gels (1 h, 140 V) after addition of Laemmle buffer devoid of DTT. After staining by overnight incubation in Instant*Blue* (Sigma Aldrich, France), gels were washed twice in ultrapure water and migration results were digitalized using a ChemiDoc XRS+ imaging system controlled by Image Lab software version 6.0.0 (BioRad, France).

### MALDI TOF mass spectrometry

MALDI TOF mass spectra on Cry11Aa were acquired on an Autoflex mass spectrometer (Bruker Daltonics, Bremen, Germany) operated in linear positive ion mode. External mass calibration of the instrument, for the *m/z* range of interest, was carried out using as calibrants the monomeric (66.4 kDa) and dimeric (132.8 kDa) ions of BSA (reference 7030, Sigma Aldrich). Just before analysis, crystals of Cry11Aa were firstly dissolved in acetonitrile/water mixture (70:30, *v/v*). For samples under reducing condition, DTT was added at a final concentration of 10 mM. The obtained solutions were therefore directly mixed in variable ratios (1:5, 1:10, 1:20, *v/v*) with sinapinic acid matrix (20 mg/mL solution in water/acetonitrile/trifluoroacetic acid, 70:30:0.1, *v/v/v*, Sigma Aldrich) to obtain the best signal-to-noise ratio for MALDI mass spectra. 1 to 2 μL of these mixtures were then deposited on the target and allowed to air dry (at room temperature and pressure). Mass spectra were acquired in the 10 to 160 kDa *m/z* range and data processed with Flexanalysis software (v.3.0, Bruker Daltonics).

MALDI TOF mass spectra on Cry11Ba were collected at the USC Mass Spectrometry Core Facility, Los Angeles, CA, USA. Purified Cry11Ba protein was dissolved in water (~ 5 mg/mL) and heated at 70 °C for 10 min to facilitate dissolution. One microliter of protein solution was spotted on a 384 Big Anchor MALDI target and let dry at room temperature. Crystallized protein was washed on-target twice with MQ water, on top of which 0.5 μL of 2,6 dihydroxyacetophenone (DHAP) solution (30 mg/ml in 50% acetonitrile:0.1% formic acid) was spotted and let dry at room temperature. Crystallized sample was then analyzed using Bruker Rapiflex^®^ MALDI-TOF MS equipped with a Smartbeam 3D, 10 kHz, 355 nm Nd:YAG laser. The laser parameters were optimized as follows: scan range = 26 μm; number of shots per sample = 1000; laser frequency = 5000 Hz. The mass spectrometer was calibrated for high-mass range using Protein A and Trypsinogen standards under Linear Mode. Data were analyzed using FlexAnalysis software and plotted using Graphpad Prism.

### In-gel digestion and peptide mass fingerprinting of Cry11Aa using MALDI

Selected bands were in-gel digested with trypsin as previously described^100^. MALDI mass spectra of the tryptic peptides were recorded on an Autoflex mass spectrometer (Bruker Daltonics, Bremen, Germany) in the reflectron positive ion mode. Before analysis samples were desalted and concentrated on RP-C18 tips (Millipore) and eluted directly with 2 μl of α-cyano-4-hydroxy cinnamic acid matrix (10 mg/ml in water/acetonitrile/trifluoroacetic acid: 50/50/0.1, *v/v/v*) on the target.

### In-gel digestion and peptide mass fingerprinting of Cry11Ba using GeLC-MS/MS

Gel Liquid Chromatography tandem mass spectrometry mass spectra collected on Cry11Ba were acquired on a ThermoFisher Q-Exactive Plus (UCLA Molecular Instrumentation Center, Los Angeles, CA, USA). Before analysis, the Cry11Ba crystals were diluted at a 1:5 dilution with ultrapure H2O and 4x SDS Loading Buffer Dye. These samples were then boiled for 3 min at 98°C and were loaded on a 4-12% Bis-Tris SDS-PAGE gel. Protein embedded in gel bands were extracted and digested with 200 ng trypsin at 37°C overnight. The digested products were extracted from the gel bands in 50% acetonitrile/49.9% H2O/ 0.1% trifluoroacetic acid (TFA) and desalted with C18 StageTips prior to analysis by tandem mass spectrometry. Peptides were injected on an EASY-Spray HPLC column (25 cm x 75 μm ID, PepMap RSLC C18, 2 μm, ThermoScientific). Tandem mass spectra were acquired in a data-dependent manner with a quadrupole orbitrap mass spectrometer (Q-Exactive Plus Thermo Fisher Scientific) interfaced to a nanoelectrospray ionization source. The raw MS/MS data were converted into MGF format by Thermo Proteome Discoverer (VER. 1.4, Thermo Scientific). The MGF files were then analyzed by a MASCOT sequence database search.

### Native mass spectrometry

Crystals of Cry11Aa were centrifuged for 5 minutes at 5000 g during the buffer wash and washed twice with ammonium acetate buffer (pH adjusted to 6.4 with acetic acid). Pelleted crystals were then dissolved in ammonium acetate buffer (pH adjusted to 11.5 using ammonium hydroxide). Gold-coated capillary emitters were prepared as previously described and used to load the protein sample ^101^. The sample was analyzed on a Synapt G1 mass spectrometer (Waters Corporation). The instrument was tuned to preserve non-covalent interactions. Briefly, the capillary voltage was set to 1.60 kV, the sampling cone voltage was 20 V, the extraction cone voltage was 5 V, the source temperature was 80 °C, the trap transfer collision energy was 10V, and the trap collision energy (CE) was set at 30 V. For MS/MS characterization, a particular charge state was isolated in the quadrupole and the complex was dissociated by application of 200V of CE. The data collected were deconvoluted and analyzed using UniDec ^102^.

### Heat stability and aggregation propensity

The thermal unfolding of Cry11Aa WT and mutants was measured by following changes as a function of temperature (15 – 95 °C) in tryptophan fluorescence leading to an increase of the F350/F330 ratio. Scattering was also monitored to address aggregation propensity of Cry11Aa WT and of the mutants F17Y, Y272Q, Y349F, Y449F, D507N-D514N and E583Q (Supplementary Fig. 12). All the measurements were performed on a Prometheus NT.48 (Nanotemper) following manufacturer’s instructions.

## Data availability

Structures and structure factor amplitudes have been deposited in the PDB databank under accession codes 7QX4 (Cry11Aa WT, pH 7.0; 10.2210/pdb7QX4/pdb), 7QX5 (Cry11Aa Y449F, pH 7.0; 10.2210/pdb7QX5/pdb), 7QX6 (Cry11Aa E583Q, pH 7.0; 10.2210/pdb7QX6/pdb), 7QX7 (Cry11Aa F17Y, pH 7.0; 10.2210/pdb7QX7/pdb), 7QYD (Cry11Ba WT, pH 6.5; 10.2210/pdb7QYD/pdb), 7R1E (Cry11Ba WT, pH 10.4; 10.2210/pdb7R1E/pdb). Raw image files are deposited in cxi.db accession number 190 (https://www.cxidb.org/id-190.html). The source data for Figure 3 and for Supplementary Figs. 5, 6, 7, 8, 10, 11 and 12 are provided in a combined Source Data file.

## Acknowledgements

We thank Aline LeRoy for help during experiments with the Nanotemper apparatus and Dr. Guy Schoehn for his support. IBS acknowledges integration into the Interdisciplinary Research Institute of Grenoble (IRIG, CEA). We thank the LCLS for beamtime allocation under proposals cxi0416, L091, P127, P125 and P141, and the EuXFEL for beamtime allocation under proposals P2156 and P2545. This work was supported by the Agence Nationale de la Recherche (grants ANR-17-CE11-0018-01 and ANR-2018-CE11-0005-02 to J.-P.C.), the CNRS (PEPS SASLELX grant to MW) and used the atomic force microscopy (AFM) platform at the IBS and the electron microscopy (EM) platform of the Grenoble Instruct-ERIC center (ISBG; UMS 3518 CNRS-CEA-UGA-EMBL) within the Grenoble Partnership for Structural Biology (PSB). Platform access was supported by FRISBI (ANR-10-INBS-05-02) and GRAL, a project of the University Grenoble Alpes graduate school (Ecoles Universitaires de Recherche) CBH-EUR-GS (ANR-17-EURE-0003). The EM facility is supported by the Auvergne-Rhône-Alpes Region, the Fondation Recherche Medicale (FRM), the fonds FEDER and the GIS-Infrastructures en Biologie Sante et Agronomie (IBiSA). Use of the LCLS at the Stanford Linear Accelerator Center (SLAC) National Laboratory, is supported by the US Department of Energy, Office of Science, and Office of Basic Energy Sciences under contract no. DE-AC02-76SF00515. The CXI instrument was funded by the Linac Coherent Light Source Ultrafast Science Instruments project, itself funded by the DOE Office of Basic Energy Sciences. Parts of the sample injector used at LCLS for this research were funded by the National Institute of Health, P41GM103393, formerly P41RR001209. This research was also supported by NIH grant GM117126. Data processing was supported by National Institutes of Health grant GM117126 to N.K.S. N.A.S.’s contributions reported in this publication were supported by the Ruth L. Kirschstein National Research Service Award of the National Institutes of Health under award number GM007185.

## Author contributions

G.T., A.-S.B, D.B., E.L., L.D., H.-W.P., B.F. designed and constructed and transformed WT and mutants plasmids; G.T., E.A.A., A.-S.B, N.A.S., D.B., N.Z., H.-W.P. and B.F. produced crystals *in vivo*; L.S., E.J.C. and M.A.B. performed MALDI-TOF mass spectrometry experiments; P.Q. and A.L. performed native MS/MS mass spectrometry experiments; G.T., E.A.A. and N.A.S. performed solubilization assays; E.A.A. performed heat stability assays; N.A.S., M. B., W.L.L. and I.G. conducted transmission electron microscopy imaging; G.T., E.A.A., A.-S.B, N.A.S., R.S. and I.S. performed crystal visualization by SEM; G.T., J.-M.T., D.F. and J.-L.P. performed crystal visualization and size measurements by AFM; G.T. and J.-L.P. performed the statistical analysis of solubilization data and G.T. and J.-L.P. performed the statistical analyses of AFM data; A.-S.B., M.B., M.W. and J.-P.C secured beamtime at the ESRF for crystal screening; M.R.S., N.S., J.R., B.F., and D.C. secured beamtime at the APS for crystal screening; M.R.S., A.-S.B., M.S.H, J.R., M.W., N.K.S., B.F., D.C., I.S., J.-P.C secured beamtime at the LCLS for data collection; I.S. and J.P.C. secured beamtime at the EuXFEL for data collection; S.E., E.G., A.R., C.C., F.R. and O.M. synthesized TbX-o4; G.T. derivatized Cry11Aa crystals for injection at LCLS; G.T., M.R.S., E.A.A., A.-S.B and N.A.S. prepared crystals for data collection at XFEL and synchrotrons; R.G.S. developed and operated the MESH-on-a-stick injector; R.L.S. and R.B.D. developed the GDVN injector; E.A.A., G.S., M.G., G.N.-K., M.K., G.S., M.S., R.L.S. and R.B.D. operated the GDVN injector; G.T., M.R.S., E.A.A., A.-S.B, N.A.S., A.S.B., M.G., G.N.-K., M.S.H., M.K., R.G.S., G.S., M.S., I.D.Y., A.G., A.B., S.B., T.M.R.S, J.R., R.L.S., R.B.D., M.W., N.K.S., D.C., I.S. and J-P.C. performed serial data collection at the LCLS; G.T., E.A.A., A.-S.B, N.C., M.G., G.N.-K., M.S.H, M.K., R.G.S., G.S., A.G., M.H., L.F., J.B., R.B., R.L., A.M., T.R.M.B, R.L.S., R.B.D., I.S. and J-P.C. performed serial data collection at the EuXFEL; E.D.Z., N.C., A.S.B., I.D.Y. and N.K.S. produced new processing tools or devices; E.D.Z, N.C., A.S.B., A.G., I.D.Y., N.K.S. and J-P.C. performed serial data processing; E.D.Z. and J.P.C phased the structural data; M.R.S, E.D.Z, N.A.S. and J-P.C. performed atomic model building, refinement and structure interpretation; G.T., M.R.S., E.D.Z and J-P.C. prepared figures and tables and wrote the manuscript with input from E.A.A., A.-S.B, N.A.S., A.S.B, M.L.G., D.B., S.E., L.S., R.S., W.L.L., J.-L.P. A.L., R.L.S., D.C. and I.S. J.-P.C. designed and coordinated the project.

