## Supplementary Information Appendix for "*De novo* determination of mosquitocidal Cry11Aa and Cry11Ba structures from naturally-occurring nanocrystals" (from another version of this preprint: version 2, 2022-01-28)

by

Guillaume Tetreau[1,18], Michael R. Sawaya[2,18], Elke De Zitter[1,18], Elena A. Andreeva[1,3,19], Anne-Sophie Banneville[1,19], Nathalie Schibrosky[2,4,19], Nicolas Coquelle[5], Aaron S. Brewster[6], Marie Luise Grünbein[3], Gabriela Nass Kovacs[3], Mark S. Hunter[7], Marco Kloos[3,8], Raymond G. Sierra[7], Giorgio Schiro[1], Pei Qiao[9], Myriam Stricker[3], Dennis Bideshi[10,11], Iris D. Young[6], Ninon Zala[1], Sylvain Engilberge[1], Alexander Gorel[3], Luca Signor[1], Jean-Marie Teulon[1], Mario Hilpert[3], Lutz Foucar[3], Johan Bielecki[8], Richard Bean[8], Romain Letrun[8], Alexander Batyuk[7], Irina Snigireva[12], Daphna Fenel[1], Robin Schubert[8], Ethan J. Canfield[13], Mario M. Alba[14], Frédéric Laporte[15], Laurence Després[15], Maria Bacia[1], Amandine Roux[16], Christian Chapelle[17], François Riobé[16], Olivier Maury[16], Wai Li Ling[1], Sébastien Boutet[7], Adrian Mancuso[8], Irina Gutsche[1], Eric Girard[1], Thomas R. M. Barends[3], Jean-Luc Pellequer[1], Hyun-Woo Park[10,11], Arthur D. Laganowsky[9], Jose Rodriguez[2,4], Manfred Burghammer[12], Robert L. Shoeman[3], R. Bruce Doak[3], Martin Weik[1], Nicholas K. Sauter[6], Brian Federici[10], Duilio Cascio[2], Ilme Schlichting[3], Jacques-Philippe Colletier[1*].

**This PDF file contains:**

- Supplementary Figures 1-13
- Supplementary Tables 1 and 2

**Affiliations:**

[1]Univ. Grenoble Alpes, CNRS, CEA, Institut de Biologie Structurale, 71 Avenue des martyrs, F-38000 Grenoble, France.

[2]UCLA-DOE Institute for Genomics and Proteomics, Department of Biological Chemistry, University of California, Los Angeles, CA 90095-1570.

[3]Max-Planck-Institut für medizinische Forschung, Jahnstrasse 29, 69120 Heidelberg, Germany.

[4]Department of Chemistry and Biochemistry, University of California, Los Angeles, CA 90095

[5]Large-Scale Structures Group, Institut Laue-Langevin, F- 38000 Grenoble, France.

[6]Molecular Biophysics and Integrated Bioimaging Division, Lawrence Berkeley National Laboratory, Berkeley, CA 94720.

[7]Linac Coherent Light Source, SLAC National Accelerator Laboratory, Menlo Park, CA 94025.

[8]European XFEL GmbH, Holzkoppel 4, 22869, Schenefeld, Germany.

[9]Department of Chemistry, Texas A&M University, College Station, TX 77845.

[10]Department of Entomology and Institute for Integrative Genome Biology, University of California, Riverside, CA 92521, USA.

[11]Department of Biological Sciences, California Baptist University, Riverside, California 92504, USA.

[12]European Synchrotron Radiation Facility (ESRF), BP 220, 38043 Grenoble, France.

[13]Mass Spectrometry Core Facility, School of Pharmacy, University of Southern California, Los Angeles, CA 90089, USA.

[14]Department of Pharmacology and Pharmaceutical Sciences, School of Pharmacy, University of Southern California, Los Angeles, CA 90089, USA.

[15]Univ. Grenoble Alpes, CNRS, LECA, F-38000, Grenoble, France.

[16]Univ. Lyon, ENS de Lyon, CNRS UMR 5182, Université Claude Bernard Lyon 1, Laboratoire de Chimie, F69342 Lyon, France.

[17]Polyvalan SARL, 15 parvis René Descartes, 69342 Lyon

[18]These authors contributed equally

[19]These authors contributed equally

[*]Correspondance : (J-P.C.)

### Supplementary Figures

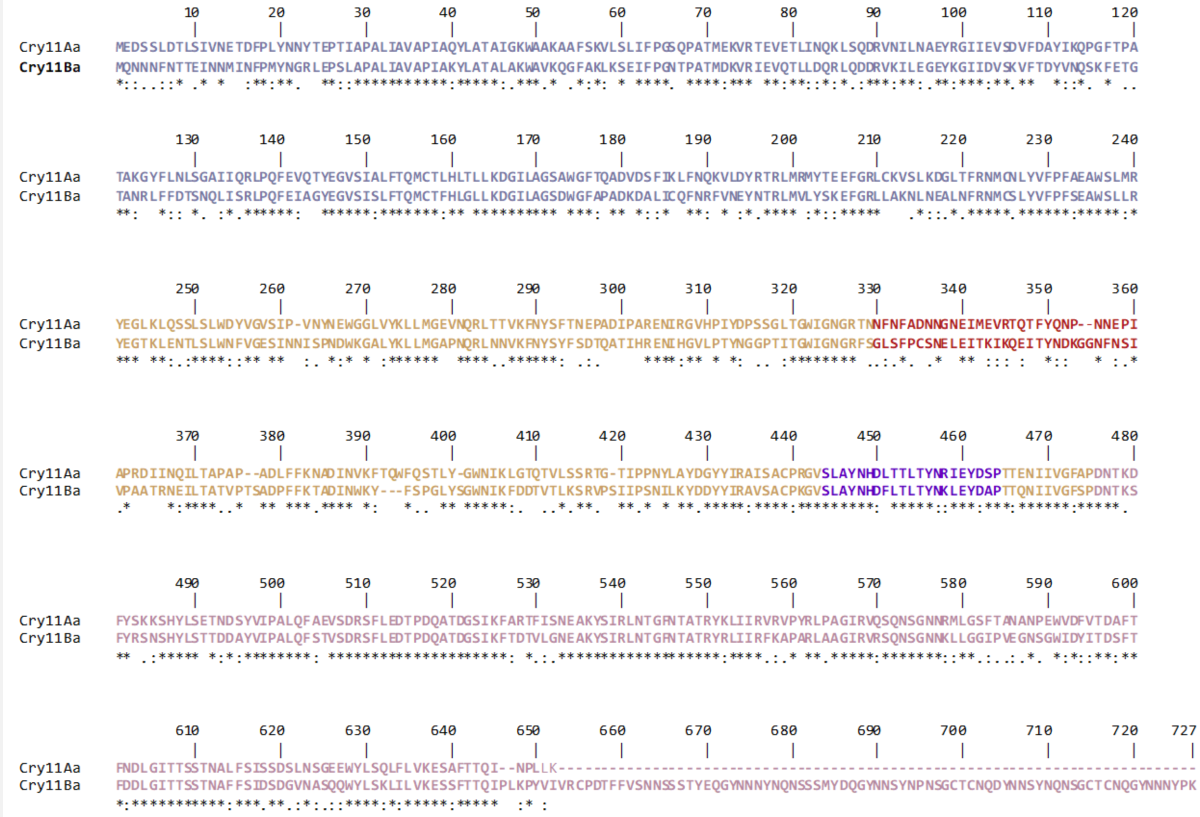

**Supplementary Fig. 1. Cry11Aa and Cry11Ba exhibit high similarity but limited sequence identity.** Cry11Aa and Cry11Ba were aligned using strap [91]. This shows that 54.0% (392) of Cry11Aa and Cry11Ba residues are identical. Domain I is shown in blue; domain II is shown in orange except for the $\alpha_h\beta_h$-handle and βpin which are shown in purple and red, respectively; domain III is shown in pink.

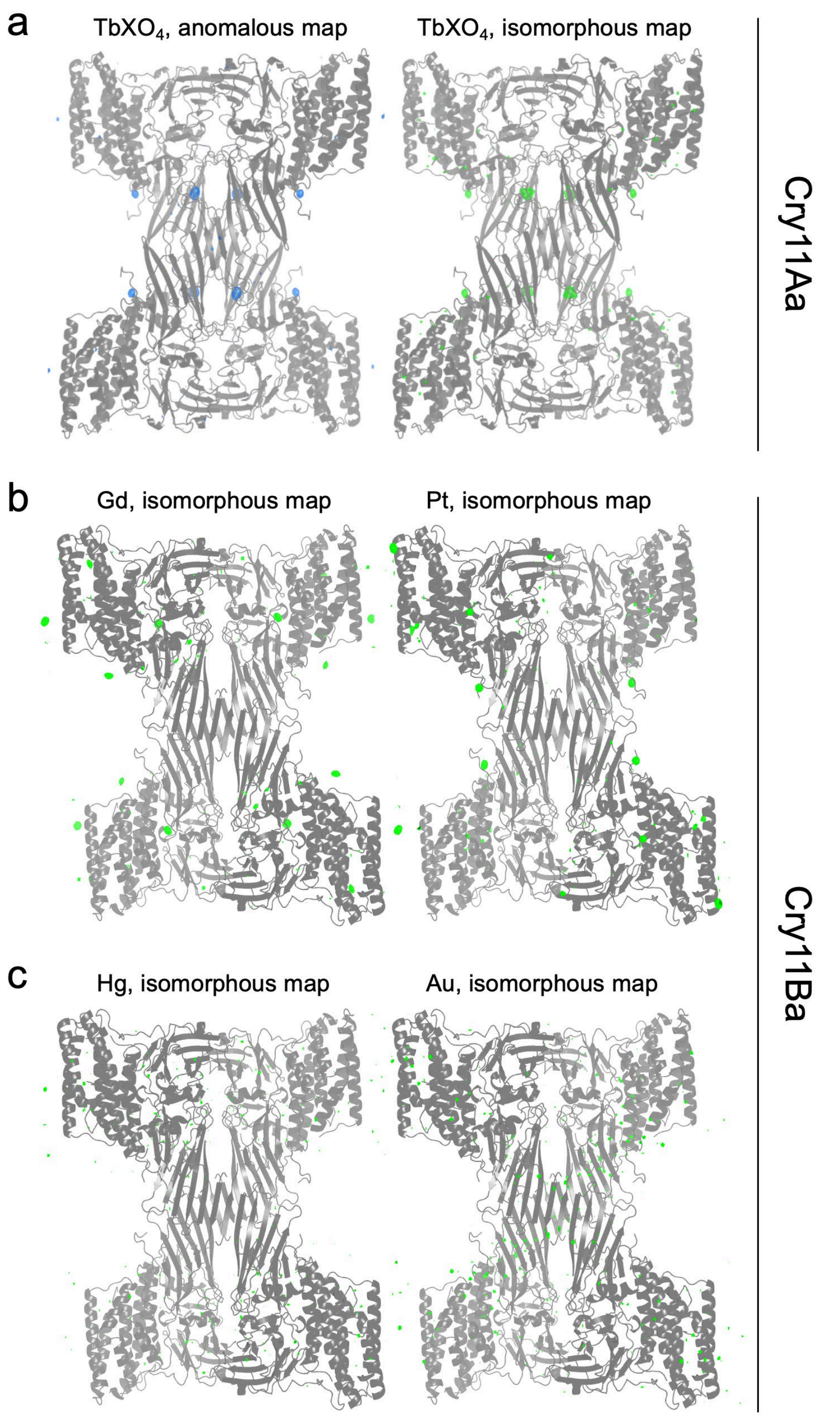


**Supplementary Fig. 2. Heavy atom locations after soaking of the Cry11 crystals. a,** Location of Tb-Xo4 in Cry11Aa is clearly indicated the anomalous (left; highest peak at 33.1 σ) and isomorphous difference maps (right; highest peak at 24.5 σ), both contoured at ± 4 σ. **b-c,** Native Cry11Ba structure revealed a posteriori that soaking of Cry11Ba with salts of gadolinium and platinum (**b**) led to their successful binding to the crystalline Cry11Ba, as revealed by isomorphous peaks > 9.3 and 7.5 σ, respectively. However, no anomalous signal was detected (highest peaks at 5.5 and 5.0 σ, respectively). Soaking of Cry11Ba crystals with salts of mercury or gold was fully infructuous, with significant peaks visible neither in the isomorphous (highest peaks at 5.3 and 5.5 σ, respectively) nor the anomalous (highest peaks at 4.9 and 5.5 σ, respectively) maps. In **b** and **c**, the isomorphous difference maps are contoured at ± 4 σ.

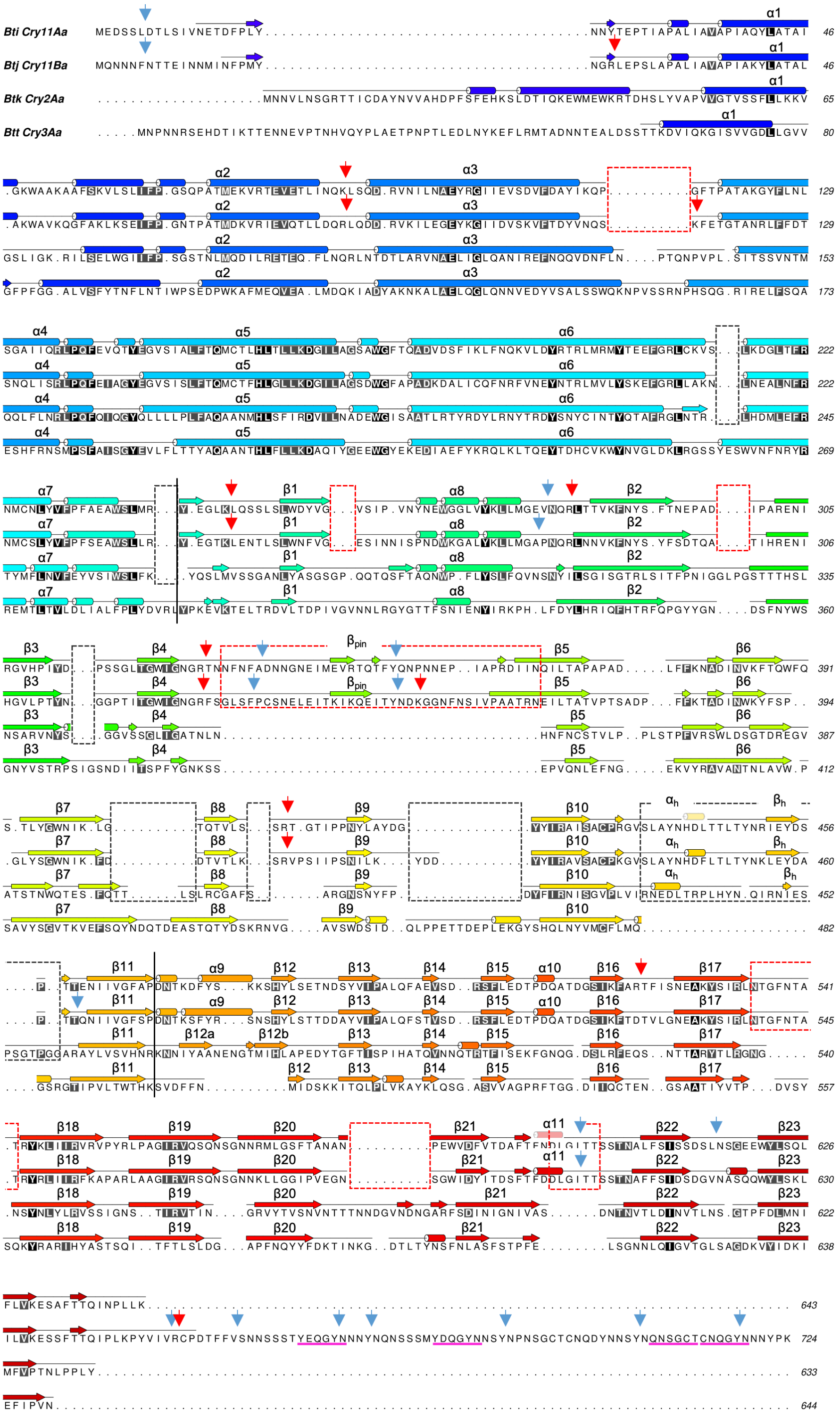


**Supplementary Fig. 3. Secondary structure assignment of Cry11Aa, Cry11Ba, Cry2Aa and Cry3Aa.** Secondary structures were assigned using DSSP[103]. α-helices and β-strand are shown by rods and arrows, respectively. Vertical black lines show the domain borders. Remarkable regions of difference between the Cry11 toxins and all other Cry toxins are indicated by red dashed boxes, whereas black boxes indicate differences between Cry11 and Cry2Aa toxin as compared to other toxins. Vertical red and blue arrows indicate trypsin and proteinase K digestion sites, respectively. Regions predicted to form short adhesive motifs of the Low Complexity, Amyloid-like Reversible Kinked Segments (LARKS) type are underlined in magenta.

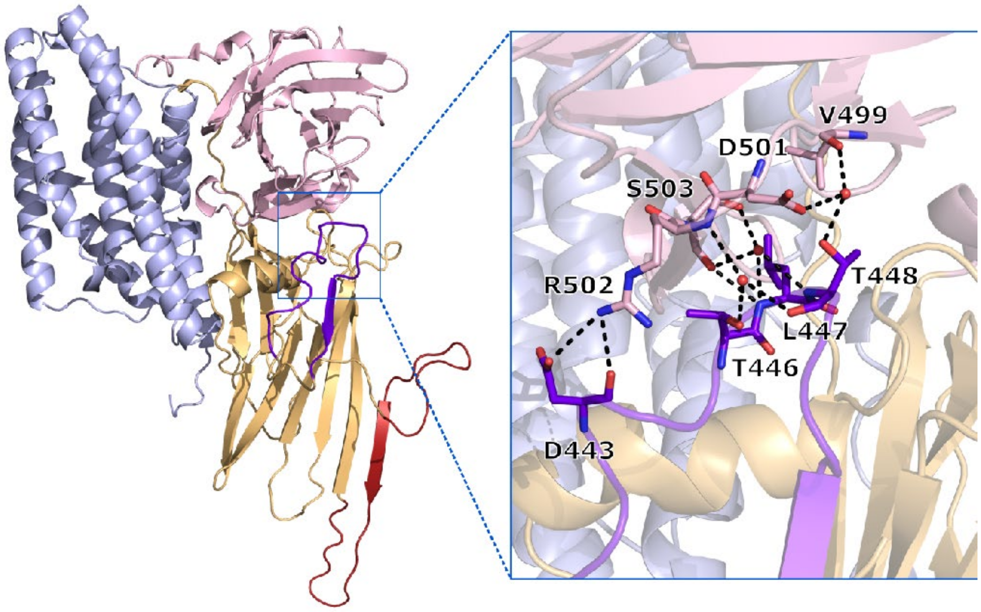


**Supplementary Fig. 4. Interactions by the $\alpha_h\beta_h$-handle in Cry11Aa.** Colour code as in Fig. 1. Dashes indicate hydrogen bonds (up to 3.2 Å) and the salt bridge between D443 and R503.

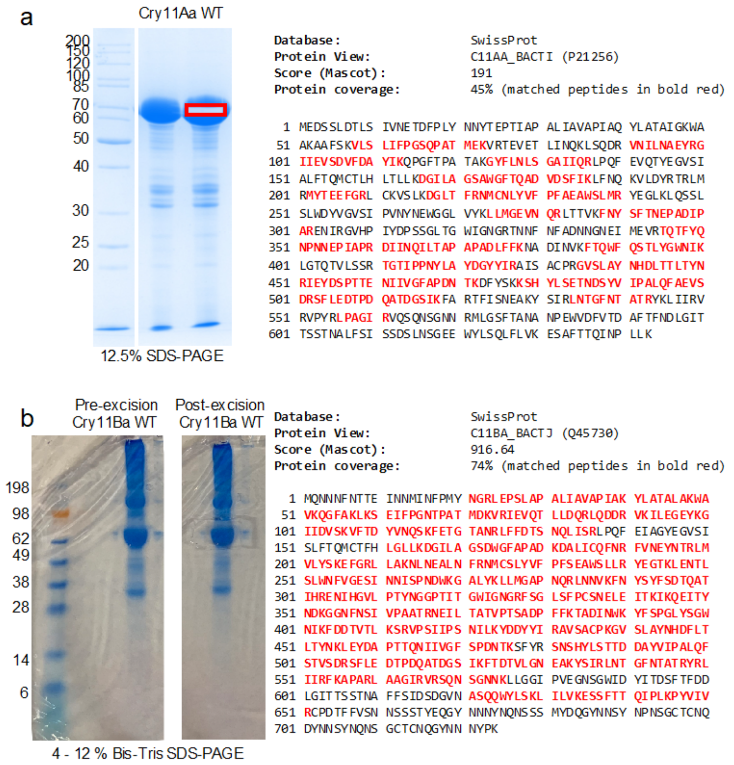


**Supplementary Figure 5. SDS PAGE and mass spectrometry confirm that Cry11Aa and Cry11Ba crystals are solely composed of these proteins. a-b,** The most abundant band present at ~70 kDa from the proteomic profile of Cry11Aa purified crystal suspension was cut and digested with protease before being analyzed in MALDI **(a)** and the most abundant band present at ~ 80 kDa was digested with trypsin before being analyzed by LC-MS/MS **(b)**. Analysis using Mascot Software allowed matching the majority of peptides identified, covering 45 and 74% of the Cry11Aa and Cry11Ba sequence, confirming that the intense band at 70 and 80 kDa are solely contributed by these proteins.

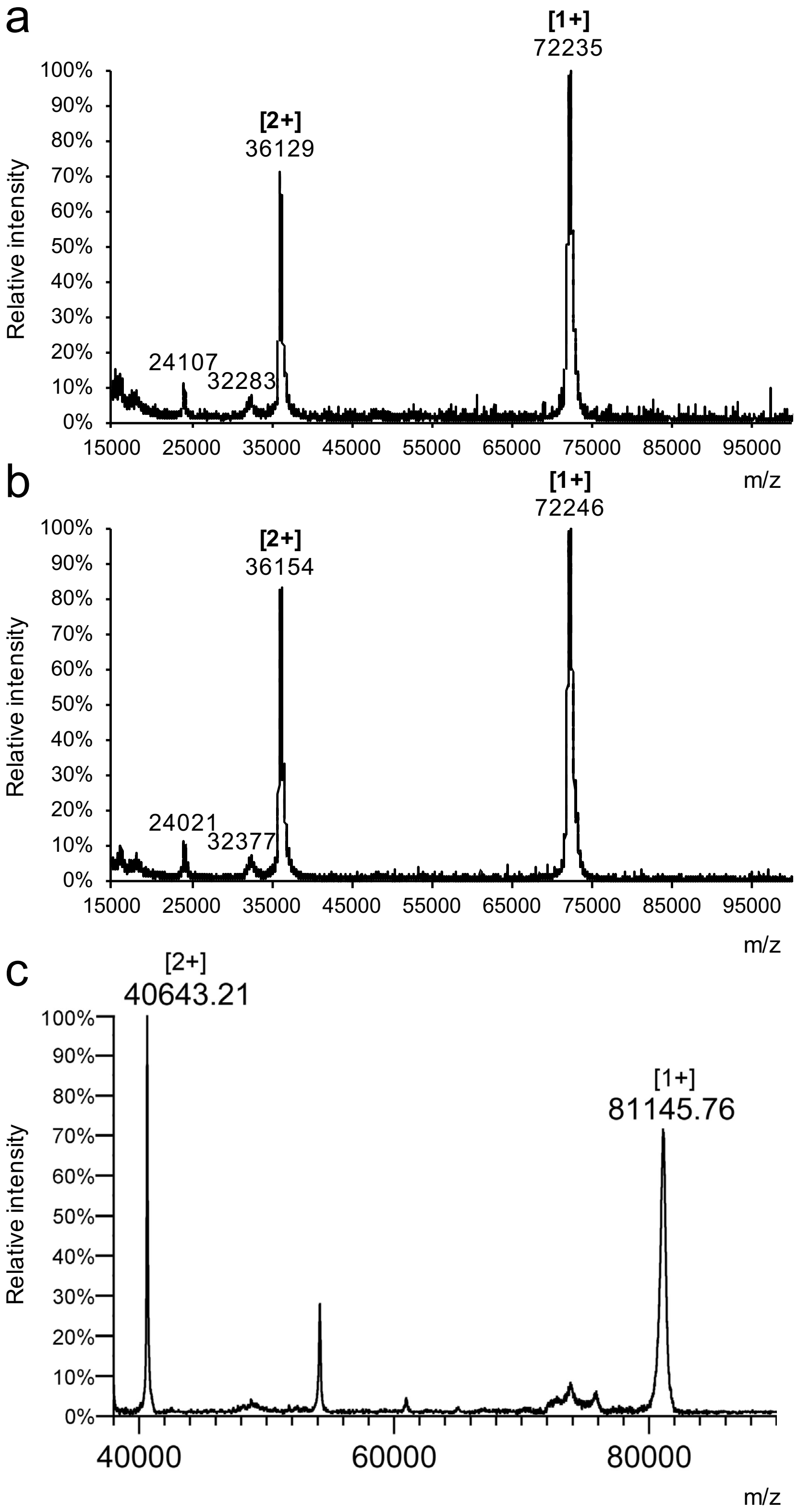


**Supplementary Fig. 6. MALDI-TOF MS analysis confirms that Cry11Aa is present as a full-size monomer in the crystal.** Crystals mixed with SA matrix in absence (**a**) or presence of 10 mM DTT (**b**) display the same profile with the most abundant peaks corresponding to the singly-charged molecular ion of a monomeric species of 72.235–72.246 kDa (mass error: ± 100 Da) (expected mass: 72.349 kDa). **c,** Cry11Ba crystals mixed with DHAP showed the presence of full-length monomer of 81.145 kDa in agreement with the predicted mass of 81.344 kDa.

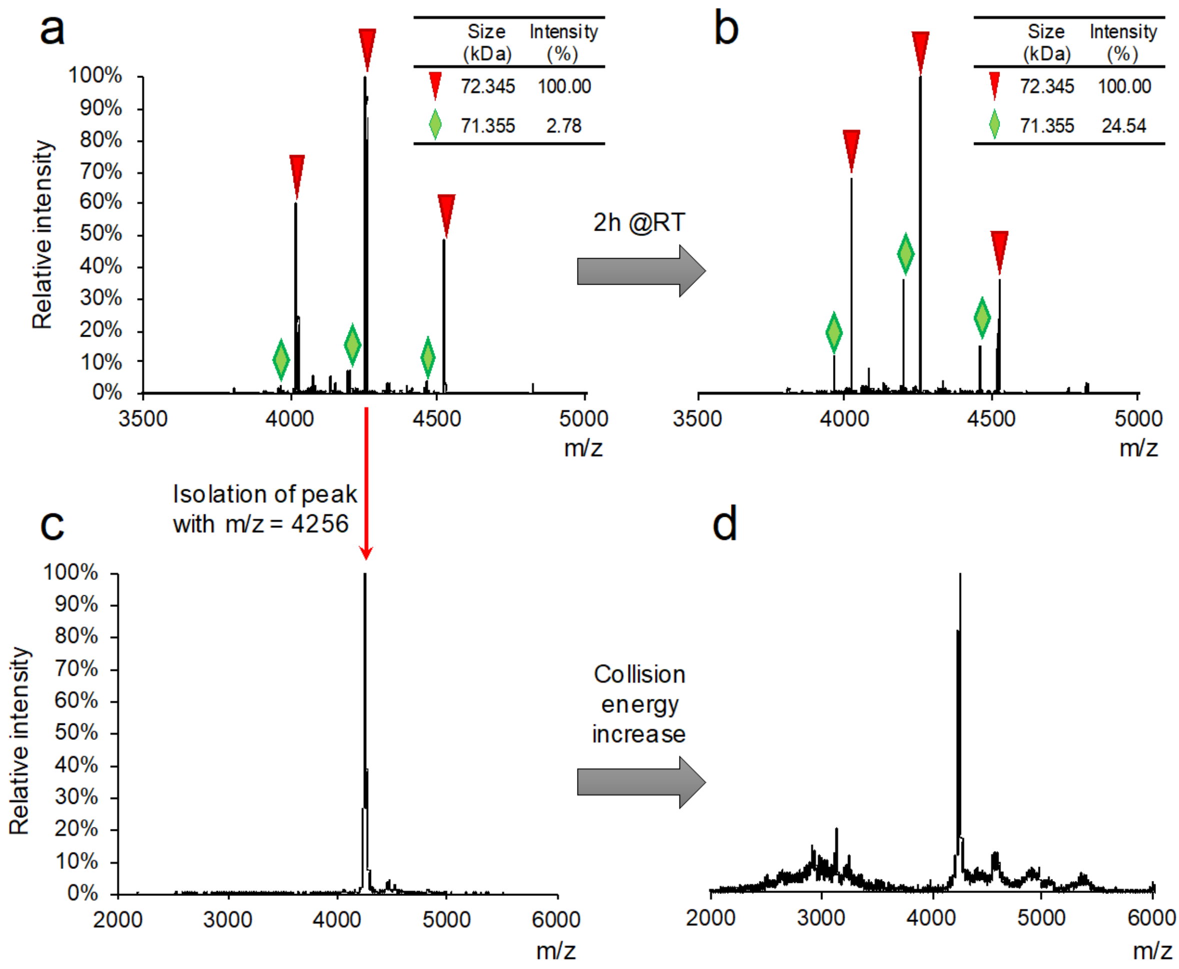


**Supplementary Fig. 7. Native mass spectrometry confirms that Cry11Aa is solubilized as a full-size monomer. a,** profile of soluble Cry11Aa in native MS shows three peaks corresponding to a full-size toxin of 72.345 kDa (expected mass: 72.349 kDa) and three peaks for a ~1 kDa smaller form much less abundant, likely resulting from a cleavage of the first 9 amino acids in N-terminal part of the toxin. **b,** incubation at room temperature leads to an increase in the smaller form, reaching one fourth of abundance of full size one, suggesting a targeted cleavage of few amino acids in the N- and/or C-terminal extremity. **c**, when isolating the most abundant peak (*i.e.,* at m/z = 4256, corresponding to Cry11Aa with a charge of 17) in the same condition as in panel a, the peak is sharp and well defined. **d,** increasing the collision energy leads to a fragmentation into species with different sizes. If the protein is an oligomer with multiple monomers through non-covalent bound, specific monomers would be easily separated and identified on the MS spectrum. This rather supports that Cry11Aa solubilizes as a full-sized monomer prone to degradation under certain conditions.

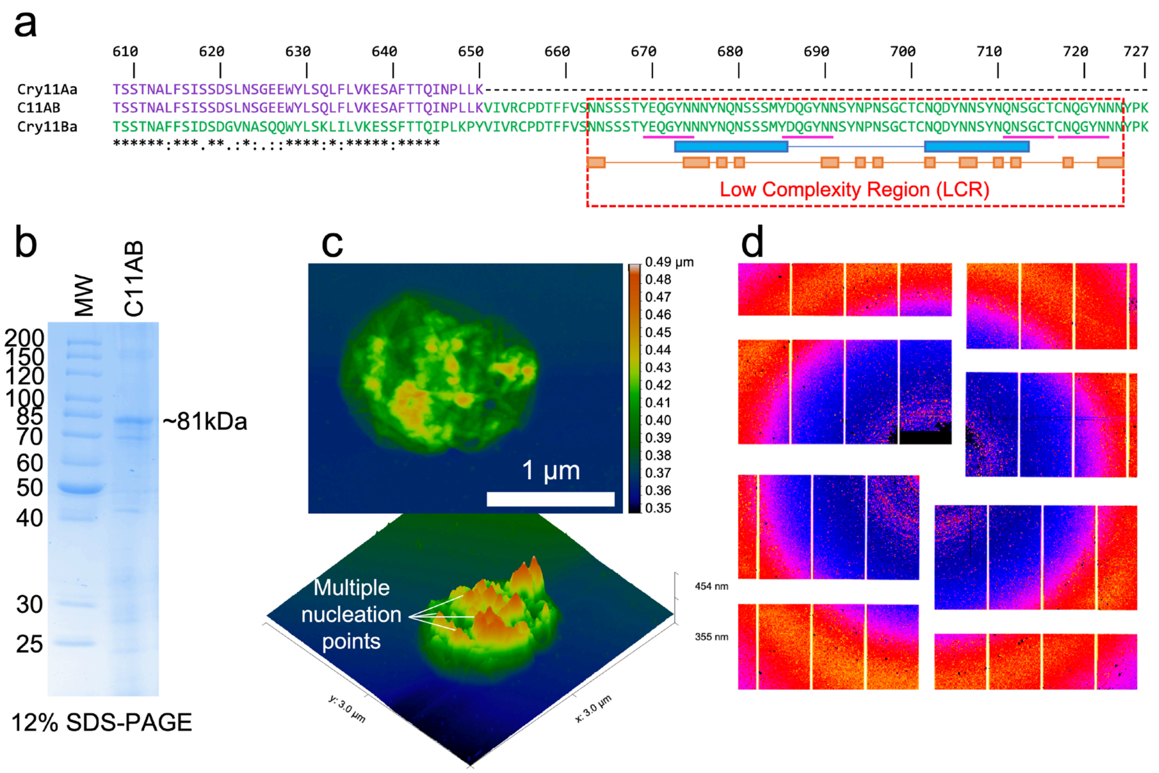


**Supplementary Fig. 8. Addition of the Cry11Ba C-terminal Low Complexity Region (LCR) to Cry11Aa does not improve its crystallization. a**, amino acid sequence alignment of Cry11Aa, Cry11Ba and Cry11Aa fused with the last 77 residues of Cry11Ba (named C11AB). Residues originating from Cry11Aa and Cry11Ba are indicated in purple and green, respectively. Blue and orange rectangles represent LCR domains identified by SEG and CAST programs, respectively. Segments predicted to form larks are underlined in magenta. **b**, C11AB crystals were purified using sucrose gradients and verified on 12% SDS-PAGE gels yielding a major band at a mean of 81.5 kDa, which matches the expected size for the fusion, although expressed at a lower level than Cry11Aa WT. **c**, purified inclusions analyzed by AFM revealed multiple nucleation points instead of a unique microcrystal. **d**, maximum projection of the few diffraction patterns that were obtained using the C11AB inclusions at the EuXFEL indicating only limited diffraction.

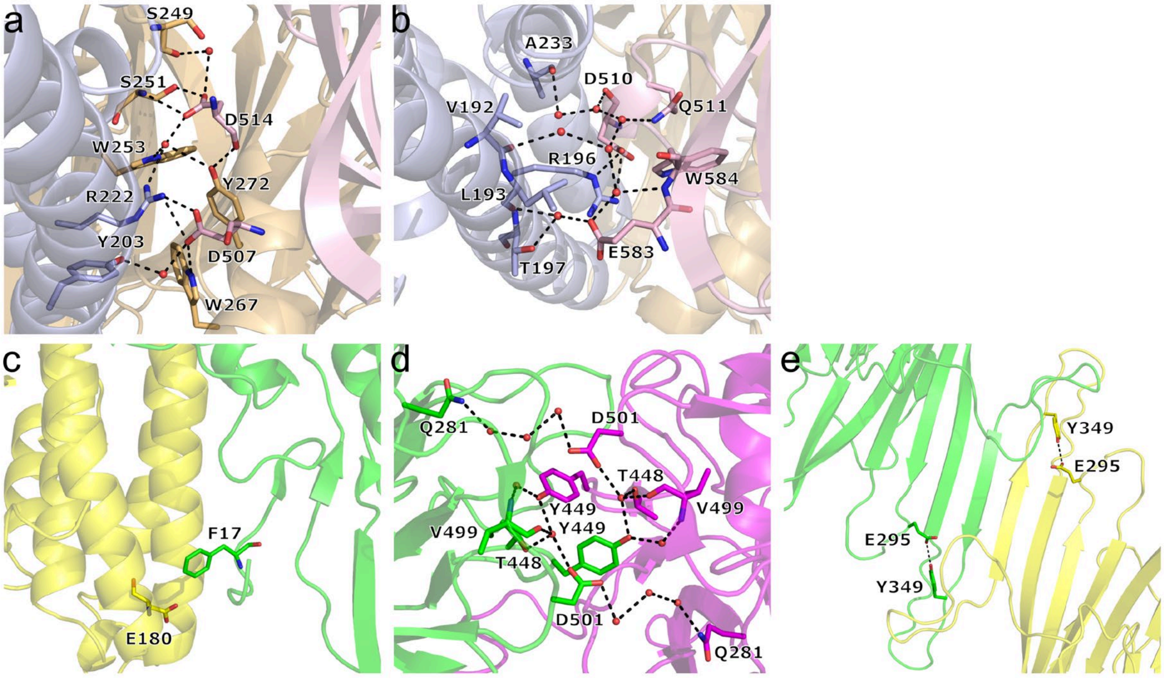


**Supplementary Fig. 9. Cry11Aa mutation strategy. a,** hydrogen and salt bonding interaction network between domain I, II and III within a single monomer formed by Y272, D504 and D514. Domain I is shown in light blue; domain II is shown in light orange; domain III is shown in pale green. For clarity, a part of domain III is not shown in cartoon mode. Hydrogen bonds are shown up until a distance of 3.2 Å. **b,** hydrogen and salt bonding interaction network between domain II and III within a single monomer involving E583. Colour code as in a. **c,** location of F17 and E80 in interface #2 between two adjacent monomers. Monomers are colour coded as in Fig. assembly. **d,** water-mediated hydrogen bonding network around Y449 in interface #1 between two adjacent monomers. For clarity, domain III of each of the monomer is omitted from the figure. Colour code as in Fig. assembly. **e,** interaction between E295 and Y349 in the $\beta_{pin}$ region which connects two adjacent monomers via interface #3. Colour code as in Fig. 2.

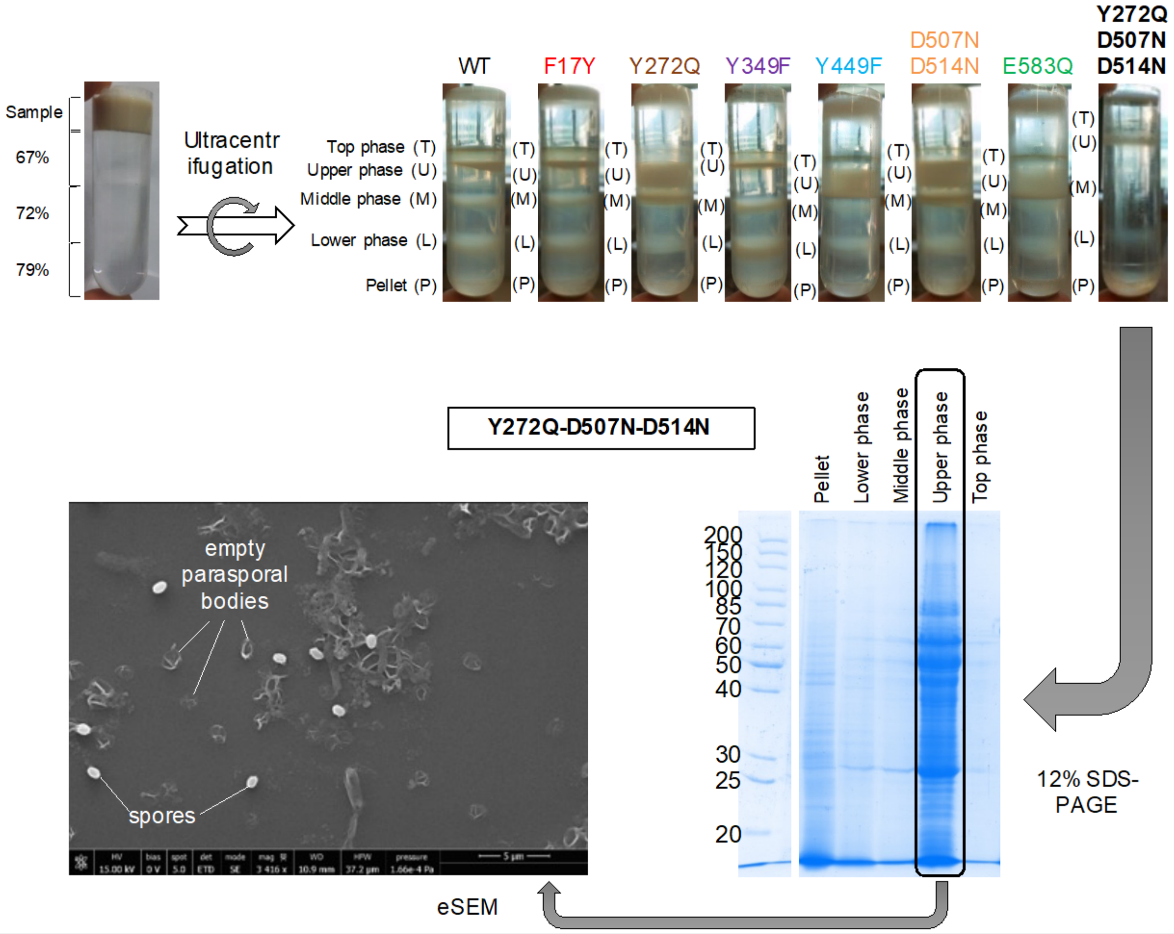


**Supplementary Fig. 10. The triple mutation Y272Q-D507N-D514N of Cry11Aa affects crystal formation.** All Cry11Aa mutants with one and two point-mutations (D507N-D514N) produced crystals that could be properly purified by sucrose gradient coupled with ultracentrifugation. In contrast, the triple mutant Y272Q-D507N-D514N showed a different purification profile. Most of proteins were contained in the upper phase of the sucrose gradient but no clear band at the ~70 kDa size could be seen. This is confirmed by environmental SEM (ESEM) experiments showing that this phase mostly contains empty parasporal bodies, suggesting that this mutation is deleterious for the proper production and/or crystallization of the toxin in the bacterium during its sporulation.

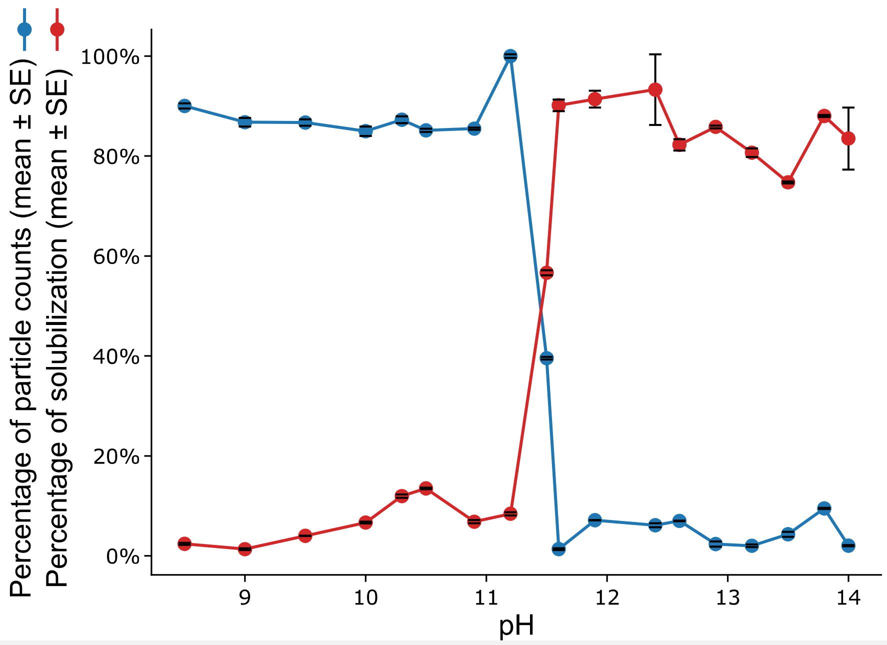


**Supplementary Fig. 11. Cry11Ba crystal solubilization as a function of pH. S**olubility and turbidity assay on Cry11Ba crystals indicate that 50 % of crystals solubilize at pH ~ 11.5. Assays were performed in triplicate. Error bars indicate the standard error of the measurements.

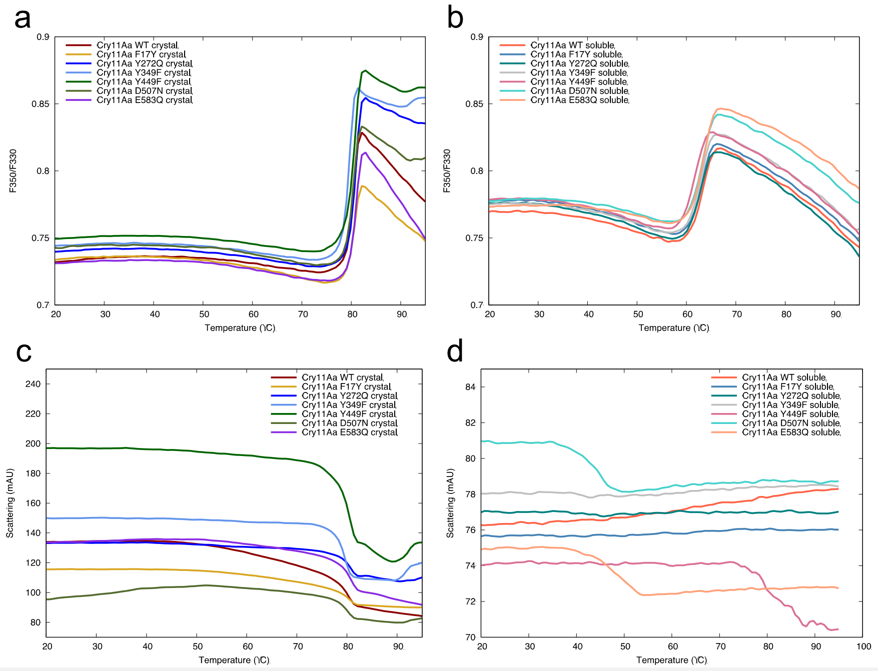


**Supplementary Figure 12. Toxin state (crystal *vs.* soluble) but not point-mutations affected the thermal stability and aggregation propensity of Cry11Aa. a-b,** differential scanning fluorometry measurements indicate the thermal unfolding of crystals (**a**) and soluble (**b**) Cry11Aa WT and mutants in function of the temperature. **c-d,** scattering measurement indicates aggregation propensity of crystals (**c**) and soluble toxins (**d**).

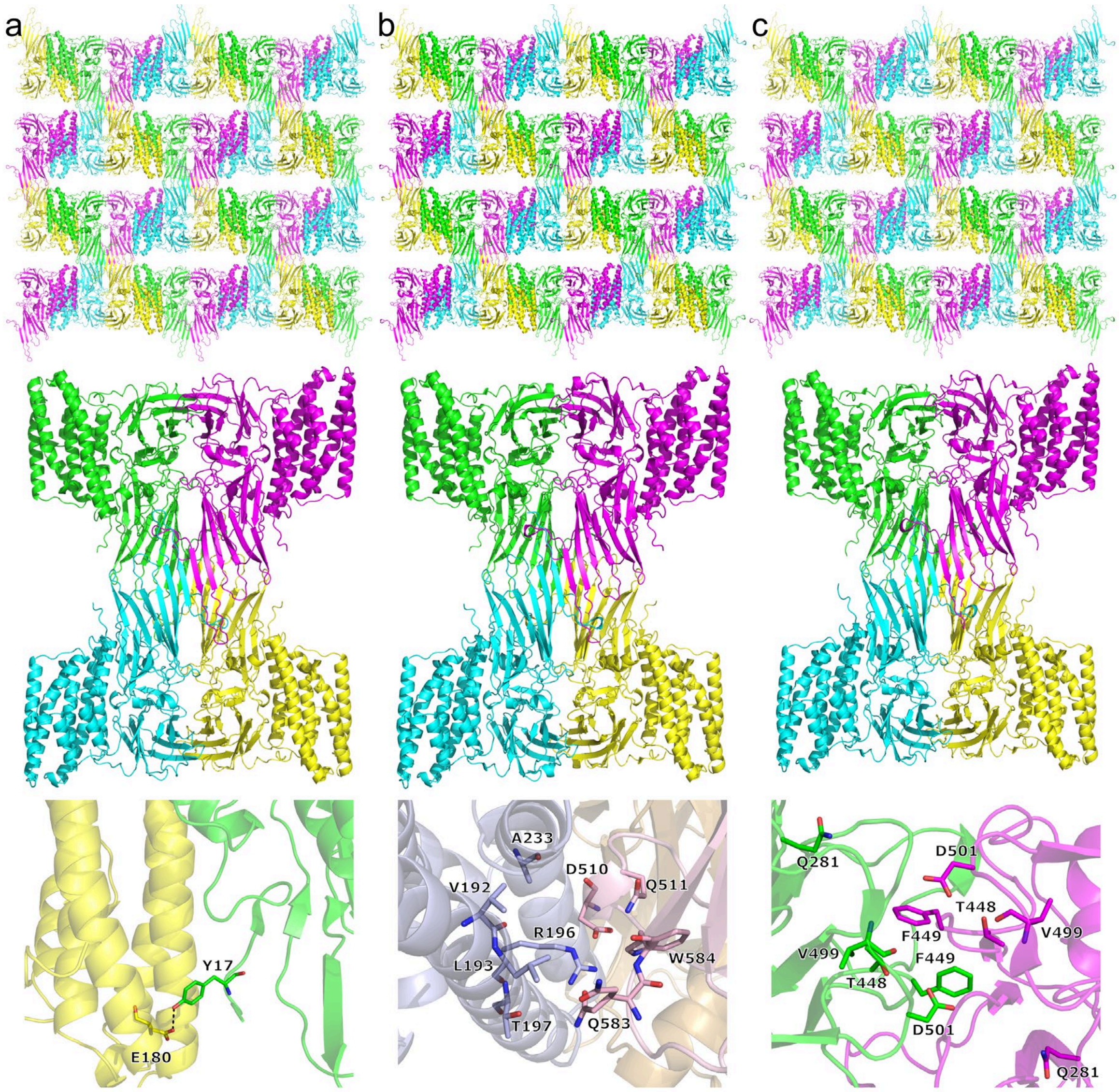


**Supplementary Fig. 13. Cry11Aa mutant crystal structures indicating the crystal packing (top), tetramer (middle) and interaction region (bottom). a,** Cry11Aa-F17Y; **b,** Cry11Aa-E583Q; **c,** Cry11Aa-Y449F. The dashes in a indicate the hydrogen bond made between E180(OE1) and Y17(OH) (2.4Å). Due to the lower resolution as compared to Cry11Aa-WT, no water molecules are observed in the interaction region. Therefore, the specific interaction pattern is not shown in **b** and **c**.

**Supplementary Table 1. Interactions in the mutated interfaces.** Hydrogen bonds up to 3.2 Å are listed.

**a. Direct and single water-mediated interactions formed by D515, Y272 and D507**

| Interaction partner 1 | Interaction partner 2 | Distance (Å)* |
|---|---|---|
| D514 (OD2) | S249 (O) | 2.9 / 2.9 |
| | S251 (OG) | 2.4 |
| D514 (OD1) | S251 (N) | 3.1 |
| | W253 (NE1) | 2.4 / 3.0 |
| | Y272 (OH) | 2.7 |
| | R222 (NH1) | 2.4 / 3.1 |
| Y272 (OH) | R222 (NH1) | 3.2 / 3.1 |
| D507 (OD1) | R222 (NH1) | 2.5 |
| D507 (OD2) | R222 (NH2) | 3.2 |
| | Y203 (OH) | 2.7 / 2.7 |
| | W267 (NE1) | 2.9 |

* Multiple distances indicate water-mediated hydrogen bonds.

**b, Extended water-mediated interactions formed by E583 and other residues involved in this network**

| Interaction partner 1 | Interaction partner 2 | Distance (Å)* |
|---|---|---|
| E583 (OE1) | T197 (OG1) | 2.8 / 2.9 |
| | L193 (O) | 2.8 / 2.8 |
| | V192 (O) | 2.6 / 2.7 / 3.0 / 3.1 |
| | A233 (O) | 2.6 / 2.7 / 2.8 / 2.9 / 2.6 / 2.9 |
| W584 (N) | V192 (O) | 3.2 / 2.7 / 3.0 / 3.1 |
| | A223 (O) | 3.2 / 2.7 / 2.8 / 2.9 / 2.6 / 2.9 |
| Q511 (NE2) | V192 (O) | 3.0 / 2.8 / 3.0 / 3.1 |
| | A223 (O) | 3.0 / 2.9 / 2.6 / 2.9 |
| D510 (O) | V192 (O) | 3.0 / 2.9 / 2.8 / 3.0 / 3.1 |
| | A223 (O) | 3.0 / 2.6 / 2.9 |
| D510 (OD2) | R196 (NE) | 3.1 |
| | R196 (NH2) | 3.3 |

* Multiple distances indicate water-mediated hydrogen bonds.

**c, Extended water-mediated interactions formed by Y449 and D501, connecting two monomers via interface #1**

| Interaction partner monomer 1 | Interaction partner monomer 2 | Distance (Å)* |
|---|---|---|
| Y449 (OH) | V499 (N) | 2.7 / 2.8 |
| | V499 (O) | 2.9 / 2.8 |
| | T448 (OG1) | 2.9 / 2.6 |
| | D501 (OD1) | 2.9 / 2.8 |
| D501 (OD2) | Q281 (NE2) | 2.8 / 2.9 / 2.6 / 3.0 |

* Multiple distances indicate water-mediated hydrogen bonds.

**d, Hydrogen bonds made by Y349**

| Interaction partner monomer 1 | Interaction partner monomer 2 | Distance (Å) |
|---|---|---|
| Y349 (OH) | E295 (OE1) | 3.1 |

**Supplementary Table 2.** Primers used to generate the mutants.

| Mutation | Forward / Reverse | Primer sequence (5'-3') | Comment |
|---|---|---|---|
| / | Forward | GCCGCAGTGTTATCACTCATGGTTATGGC | Amp_F1 was used with each reverse primer of Cry11Aa mutant construction. For each mutation primer, the inserted mutation is indicated as a capital bold letter. To generate the vector containing the mutation, the two fragments generated for the same mutations were assembled by Gibson assembly following the procedure described in the manuscript. |
| F17Y | Reverse | TTATATAATGGA**T**AGTCTGTTTCATTAACTATACTTAAAGTATCTAAAGAACTATCT | |
| Y272Q | Reverse | TAATAACTT**T**T**G**AACTAGTCCTCCCCATTCATTATAATTTACAGG | |
| Y349F | Reverse | TGGATTTTGA**A**AAAAAGTTTGTGTTCTAACTTCCATAATTT | |
| Y449F | Reverse | TCTATTCTATTA**A**ATGTTAGTGTTGTAAGATCGTGATTATATGCAAG | |
| D507N-D514N | Reverse | TTGCTTGATCTGGCGTAT**T**TTCTAAAAATGATCTATCT | |
| E583Q | Reverse | ATCCACCCATT**G**TGGATTAGCATTTG | |
| / | Reverse | AGTGCTGCCATAACCATGAGTGATAACACT | Amp_R1 was used with each forward primer of Cry11Aa mutant construction. For each mutation primer, the inserted mutation is indicated as a capital bold letter. To generate the vector containing the mutation, the two fragments generated for the same mutations were assembled by Gibson assembly following the procedure described in the manuscript. |
| F17Y | Forward | GTTAATGAAACAGACT**A**TCCATTATATAATAATTATACCGAACCTACT | |
| Y272Q | Forward | GGGAGGACTAGTT**C**A**A**AAGTTATTAATGGGGGA | |
| Y349F | Forward | CACAAACTTTTT**T**TCAAAATCCAAATAATGAGCCT | |
| Y449F | Forward | TTACAACACTAACAT**T**TAATAGAATAGAGTATGATTCACCTACTACAG | |
| D507N-D514N | Forward | ATACGCCAGATCAAGCAACA**A**ACGGCAGTATTAAATTTG | |
| E583Q | Forward | TAATCCA**C**AATGGGTGGATTTTGTCACAG | |
| C11AB chimera | Forward | CTATCCTAAATAGGCGATCGCACTCATTAGGC | This fragment was obtained using Amp_R1 as reverse primer. |
| C11AB chimera | Reverse | CGTACAATAACCTTTAGTAACGGATTAATTTGCGTCGTAAAGG | This fragment was obtained using Amp_F1 as forward primer. |
| C11AB chimera | Forward | CGTTACTAAAGGTTATTGTACGTTGTCCGGATACTTTTTTTGTG | This fragment was assembled with the two abovementioned fragments by Gibson to create the shuffle vector encoding the C11AB chimera. |
| C11AB chimera | Reverse | CGATCGCCTATTTAGGATAGTTATTGTTATACCCTTGGTTACATGTACAG | |
